# A native single-transcript TnpB architecture enables efficient virus-induced genome editing and visual screening of heritable progeny harboring phenotypically silent edits

**DOI:** 10.64898/2026.09.09.750255

**Authors:** Rui Wang, Hongfei Liu, Shuang Jia, Shasha Bai, Xingyu Cao, Dongming Li, Yongwei Sun

## Abstract

TnpB are ultra-compact RNA-guided nucleases, yet most reported systems require separate expression of the nuclease and cognate reRNA, eroding their size advantage and restricting viral delivery in plants. Here we demonstrate that ISDra2 TnpB harnesses its native overlapping architecture in which the reRNA sequence resides within the 3’ coding region of TnpB, processing its own mRNA to generate functional reRNA and enabling single-transcript genome editing in plants. Only a 7-bp conserved motif followed by a 16-20 bp spacer directly downstream of the coding sequence suffices for editing, requiring no exogenous ribozymes or independent guide promoters. This ultra-compact design enables simultaneous packaging of the editing system and an *NbPDS* silencing module into a single TRV vector, achieving robust editing in systemic *Nicotiana benthamiana* tissues and establishing a dual VIGE-VIGS strategy for direct visual selection of heritable, transgene-free edited progeny via distinct albino phenotypes, reducing laborious large-scale genotyping even for targets lacking inherent visible traits.

## Introduction

TnpB proteins derived from the IS200/IS605 transposon family are widely regarded as evolutionary ancestors of Class 2 type V CRISPR effectors, including Cas12 nucleases (Altae-Tran et al., 2021; Altae-Tran et al., 2023; Karvelis et al., 2021; Shmakov et al., 2017). Recent studies have established TnpB (∼400 amino acids) as a compact RNA-guided DNA endonuclease that associates with the right-end element RNA (reRNA) to mediate transposon-associated motif (TAM) dependent double-stranded DNA cleavage. Its minimal size compared with conventional CRISPR-Cas nucleases makes TnpB particularly attractive for applications with limited cargo capacity, such as adeno-associated virus (AAV)-mediated in vivo gene therapy (Jiang et al., 2023) and virus-induced genome editing (VIGE) in plants (Ji et al., 2025; Thornton et al., 2026). Previous studies showing that reRNA overlaps the 3’ region of tnpB genes in bacteria and archaea (Xiang et al., 2024; Xu et al., 2023). Although biochemical studies have demonstrated that TnpB can process its own transcript to generate reRNA molecules, this native overlapping configuration has not been shown to support detectable genome editing activity in *Streptomyces* (Luo et al., 2025). Consequently, reported TnpB-based genome editing systems have primarily adopted separate expression cassettes for TnpB and reRNA, thereby compromising the inherent size advantage of this compact architecture (Altae-Tran et al., 2023; Karvelis et al., 2021; Li et al., 2024; Lv et al., 2024; Molla et al., 2024; Nagalakshmi et al., 2026; Nakagawa et al., 2023; Thornton et al., 2026; Wang et al., 2024; Xiang et al., 2024). Whether this conserved single-transcript architecture can be functionally harnessed for genome editing in eukaryotic organisms, particularly plants, remains unknown.

The VIGE system bypasses tissue culture by delivering genome editing reagents directly to plant meristems, thereby eliminating somaclonal variation, reducing genotype dependence, and facilitating transgene-free, scalable genome engineering across diverse plant species (Qiao et al., 2026; Steinberger and Voytas, 2025). Although VIGE has achieved substantial progress in plants (Kang et al., 2025; Kim et al., 2023; Mikhaylova, 2025; Qiao et al., 2026; Qiao et al., 2025; Steinberger and Voytas, 2025), most reported applications have focused on the knockout of genes associated with readily observable phenotypes, such as *ChlH* or *PDS*. Applications involving genes that do not produce obvious visible phenotypes after editing remain relatively limited, largely because VIGE workflows require multiple experimental steps and extended periods for progeny generation. Moreover, owing to the spatially heterogeneous nature of viral movement and genome editing activity, editing outcomes can vary substantially among different tissues or even among individual seed capsules within the same plant, making the identification of heritable edited individuals highly laborious and often dependent on large-scale genotyping.

In this study, we demonstrate that ISDra2 TnpB can harness its native overlapping architecture, in which the reRNA sequence is embedded within the 3′ coding region of TnpB, to process its own transcript and generate functional reRNA, thereby enabling single-transcript genome editing in plants. Building on this native compact architecture, we further developed a dual VIGE–VIGS strategy by integrating an *NbPDS* silencing module into a single TRV vector, achieving robust genome editing in systemic *Nicotiana benthamiana* tissues and enabling visual identification of heritable, transgene-free edited progeny through distinct albino phenotypes. This approach provides a practical solution for reducing laborious genotyping, particularly for target genes lacking visible phenotypes.

## Results

### The native single-transcript TnpB architecture enables genome editing in rice

Among TnpB nucleases, *ISDra2* TnpB is one of the most extensively characterized systems. Previous studies have revealed that its reRNA sequence naturally overlaps with the 3’ coding region of the *ISDra2 TnpB* gene in prokaryotic loci, forming an evolutionarily conserved compact single-transcript architecture (Fig. 1a). To determine whether this native architecture can be functionally exploited for genome editing in plants, we first generated a rice codon-optimized version of the wild-type *ISDra2 TnpB* gene. Codon optimization was applied upstream of the overlapping region, while the coding sequence corresponding to the reRNA-overlapping region (224 bp) was left unoptimized. In addition, four previously reported amino acid substitutions that enhance genome editing activity were introduced (Thornton et al., 2026), generating the optimized variant designated rOpt-eTnpBc (Fig. 1b). Based on this design, we constructed two expression configurations: a single-transcript unit (STU) and a dual-transcript unit (DTU). In the STU configuration, the *ISDra2 TnpB* coding sequence was directly followed by a 7 bp motif (GGTTCAA) and a 16–20 bp spacer sequence, forming a single transcript driven by the maize *ubiquitin* promoter (Fig. 1a, b). In the DTU configuration, *ISDra2 TnpB* expression was driven by the maize *ubiquitin* promoter, while reRNA was expressed independently under the *OsU3* promoter (Fig. 1b). We generated eight constructs targeting seven endogenous rice genes, including *OsPDS*, *OsHMBPP*, *OsYSA*, *OsWaxy*, *OsCKX2*, *OsCERK1*, and *OsPi21*, and evaluated genome editing efficiencies in stable transgenic rice lines. Strikingly, the STU achieved editing efficiencies comparable to those of the DTU (Fig. 1c). Moreover, the STU system generated biallelic mutants displaying albino phenotypes in *OsHMBPP* edited rice plants (Fig. 1d). Across the eight tested target sites, no significant difference was observed at four loci (*OsHMBPP*, *OsPDS-T1*, *OsCKX2*, and *OsCERK1*), while two loci showed reduced activity (*OsYSA* and *OsPDS-T2*)and two exhibited significantly higher editing efficiency (*OsWaxy and OsPi21*) compared with the DTU system (Fig. 1c).

**Fig. 1.**
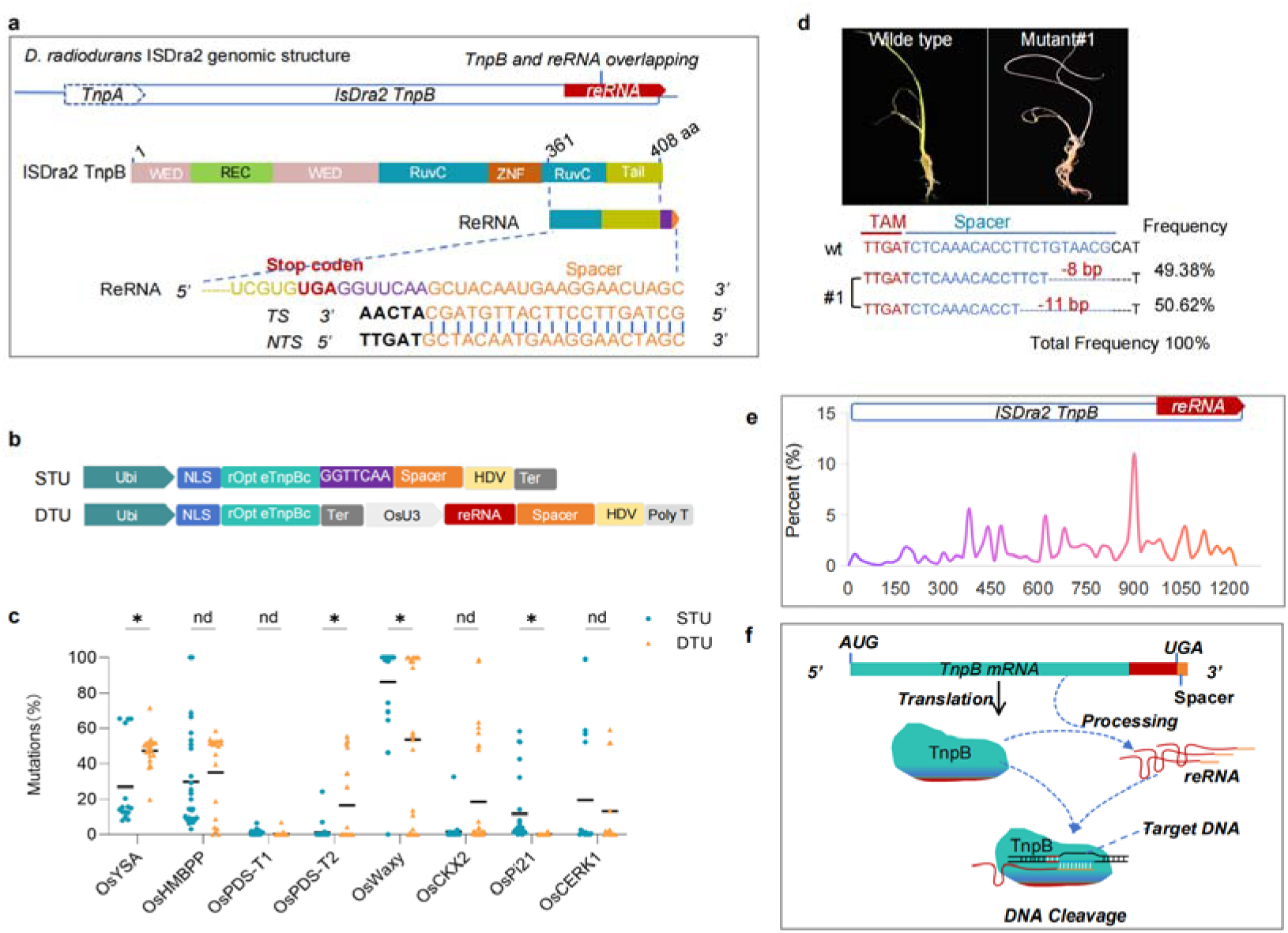
STU architecture enables genome editing in rice. **a,** Schematic representation of the *D. radiodurans* ISDra2 genomic organization and the overlapping region between *TnpB* and reRNA. **b,** Schematic diagrams of the STU and DTU constructs used for genome editing in rice. **c,** Comparison of genome editing efficiencies between STU and DTU constructs across eight endogenous target sites in rice. Data were analysed using an unpaired t-test with a two-tailed Pvalue. nd, P > 0.05; *P < 0.05; Each point represents an independent transgenic rice line (n > 15). **d**, Genotypes and albino phenotypes of biallelic *OsHMBPP* mutants generated by the STU-mediated genome editing system. **e,** Nanopore-based full-length transcript sequencing of STU-expressing transgenic rice plants. **f,** Proposed model illustrating how the STU system enables coordinated transcription, RNA processing, TnpB translation, and genome editing activity.

### A native STU architecture is conserved across TnpB orthologs

To elucidate the molecular basis underlying the STU architecture, we performed Nanopore full-length transcript sequencing of transgenic rice plants. In both STU- and DTU-expressing plants, we identified a series of 5’-truncated ISDra2 TnpB transcripts corresponding to processed reRNA-like molecules (Fig. 1e), providing direct transcriptomic evidence that ISDra2 TnpB transcripts can serve as precursors for guide RNA generation in planta. To determine whether this architecture could be extended to other TnpB orthologs, we analyzed ISAam1 TnpB, which also contains overlapping coding and reRNA regions (Supplementary Fig. 1a). Using the same single-transcript design, we validated genome editing activity in soybean hairy roots. Consistent with the results obtained with ISDra2 TnpB, the STU ISAam1 TnpB system achieved editing efficiencies comparable to those of the corresponding DTU configuration (Supplementary Fig. 1b). Together, these results demonstrate that the STU architecture is not restricted to a single TnpB ortholog and can be applied to diverse TnpB systems for plant genome engineering applications .

### The native STU architecture enables efficient VIGE in *N. benthamiana*

To investigate whether the STU architecture could be adapted for VIGE and overcome the cargo constraints associated with viral vectors. We introduced the STU cassette into the TRV2 component of a Tobacco rattle virus (TRV)-based VIGE system, generating the TRV-STU.1 vector (Fig. 2a). The *NbChlH* gene was selected as the target locus because its knockout results in a distinct chlorotic phenotype (Fig. 2b), providing a convenient visual readout of editing activity. Amplicon deep sequencing revealed high genome editing frequencies in both infiltrated leaves and systemically infected tissues, with editing frequencies reaching up to 86.3% in chlorotic systemic leaves (Fig. 2c-d). A recent study has reported that codon optimization significantly enhances TnpB-mediated genome editing efficiency in plants, with a human- and yeast-optimized variant showing superior performance in viral delivery systems (Nagalakshmi et al., 2026). Research also demonstrated that truncated reRNA (116 nt) retains substantial activity (Thornton et al., 2026). Based on these findings, we further investigated whether similar optimization strategies could enhance the performance of our STU system in VIGE. We generated two constructs, TRV-STU.2 and TRV-STU.3. In TRV-STU.2, the full-length 224-bp reRNA region was preserved without optimization, whereas TRV-STU.3 contained only the unoptimized 116-bp truncated reRNA core sequence (Fig. 2a). To evaluate whether STU provides advantages in VIGE compared with separate expression systems, we constructed an additional vector, TRV-DTU.1, in which the full-length eTnpBc and *NbChlH*-targeting reRNA were linked within a single transcript using a hammerhead (HH) ribozyme. These constructs were then introduced into the TRV2 vector and used to inoculate *N. benthamiana*. Amplicon deep sequencing results showed that both TRV-STU.2 and TRV-STU.3 further improved genome editing efficiency (Fig. 2b-e), which was consistent with the earlier onset of chlorotic symptoms and increased proportion of chlorotic systemic leaves (Fig. 2c-e). To further investigate the RNA processing characteristics of the STU architecture in the VIGE system, we performed Nanopore full-length transcript sequencing of systemic leaves infected with TRV-STU.2 and TRV-STU.3. Similar to the observations in transgenic rice, a series of continuous 5′-truncated ISDra2 TnpB transcripts were detected in systemic leaves. Notably, despite being present within the same viral transcripts, sequences encoding the TRV coat protein (CP) and other viral regions showed few detectable cleavage events. These results suggest that ISDra2 TnpB-mediated RNA processing is highly localized to the TnpB region, with limited cleavage activity toward adjacent non-TnpB sequences, supporting the localized nature of this self-processing activity (Fig. 2f-g) .

**Fig. 2.**
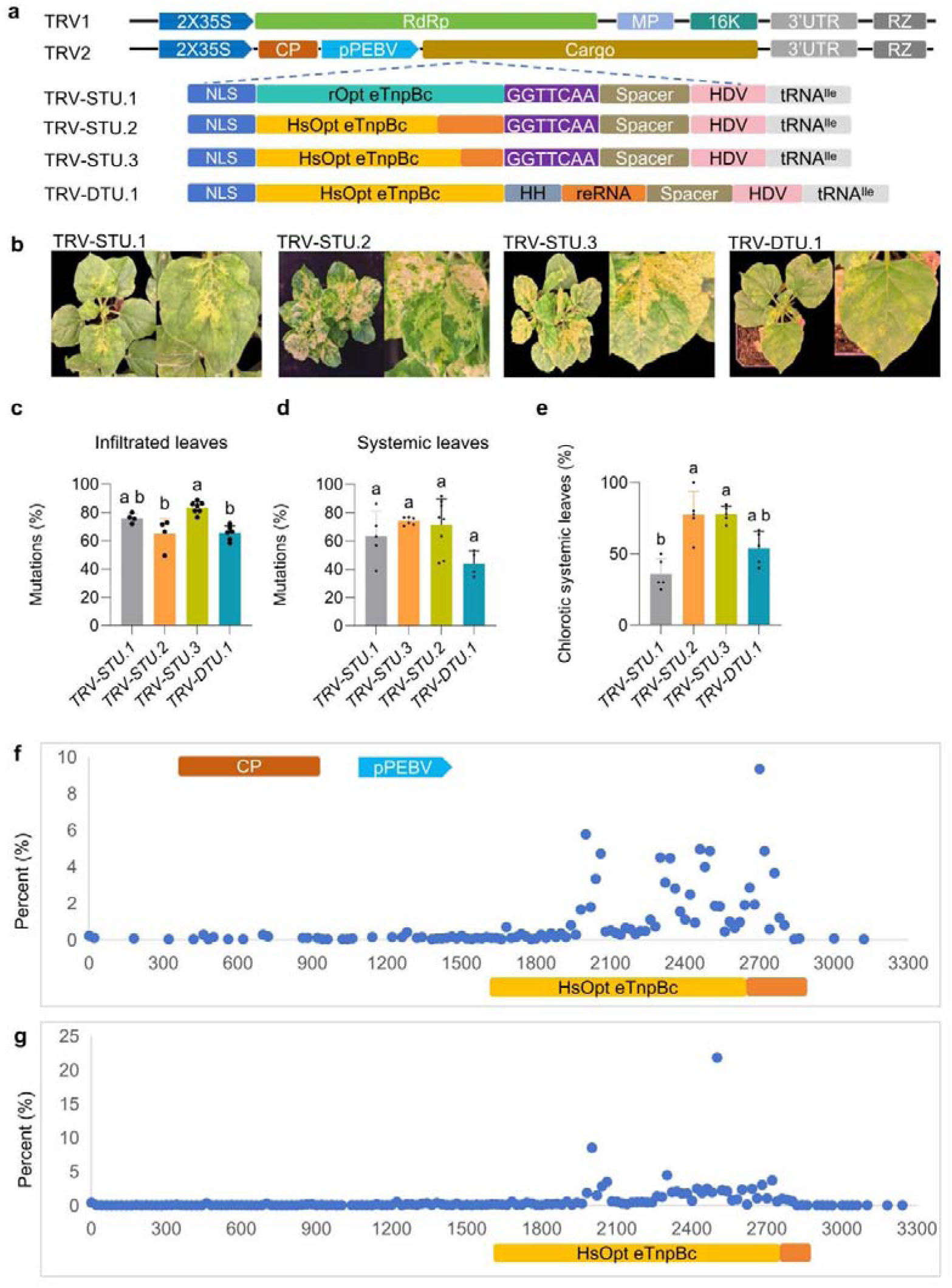
STU architecture enables genome editing in *N. benthamiana*. **a,** Schematic representation of the TRV1 and TRV2 vector systems carrying STU- or DTU-based TnpB expression configurations. **b,** Phenotypes of *N. benthamiana* plants infected with TRV vectors carrying four different TnpB configurations. Images were taken approximately 2 weeks after TRV infiltration. **c–e,** Comparison of genome editing efficiencies among four configurations in infiltrated leaves **(c)**, systemic leaves **(d)**, and the proportion of chlorotic systemic leaves **(e)**. Data are presented as mean ± standard error of the mean (SEM). One-way analysis of variance (ANOVA) followed by Tukey’s multiple comparisons test was performed to evaluate statistical differences among groups. Bars sharing any common letter are not significantly different (P > 0.05); bars without shared letters differ significantly (P < 0.05). **f-g,** Distribution and abundance of 5’ truncation sites detected within full-length TRV2 transcripts in systemic leaves infected with TRV-STU.2 (**f**) and TRV-STU.3 (**g**) as revealed by Nanopore-based full-length transcript sequencing.

To determine whether these observations corresponded to heritable genome editing, we collected seeds from chlorotic capsules and analyzed the genotypes of the resulting progeny (Fig. 3a-d). For TRV-STU.1, approximately 10%-40% of progeny seedlings exhibited a chlorotic phenotype, indicative of tetra-allelic mutations, while most green seedlings still carried editing efficiencies of ∼50% (Fig. 3e, f). In contrast, TRV-STU.2 and TRV-STU.3 showed a markedly increased proportion of chlorotic seedlings, reaching up to 100% in fully chlorotic capsules (Fig. 3e, f). TRV-DTU.1 produced approximately 30%-40% chlorotic progeny seedlings (Fig. 3e, f).

**Fig. 3.**
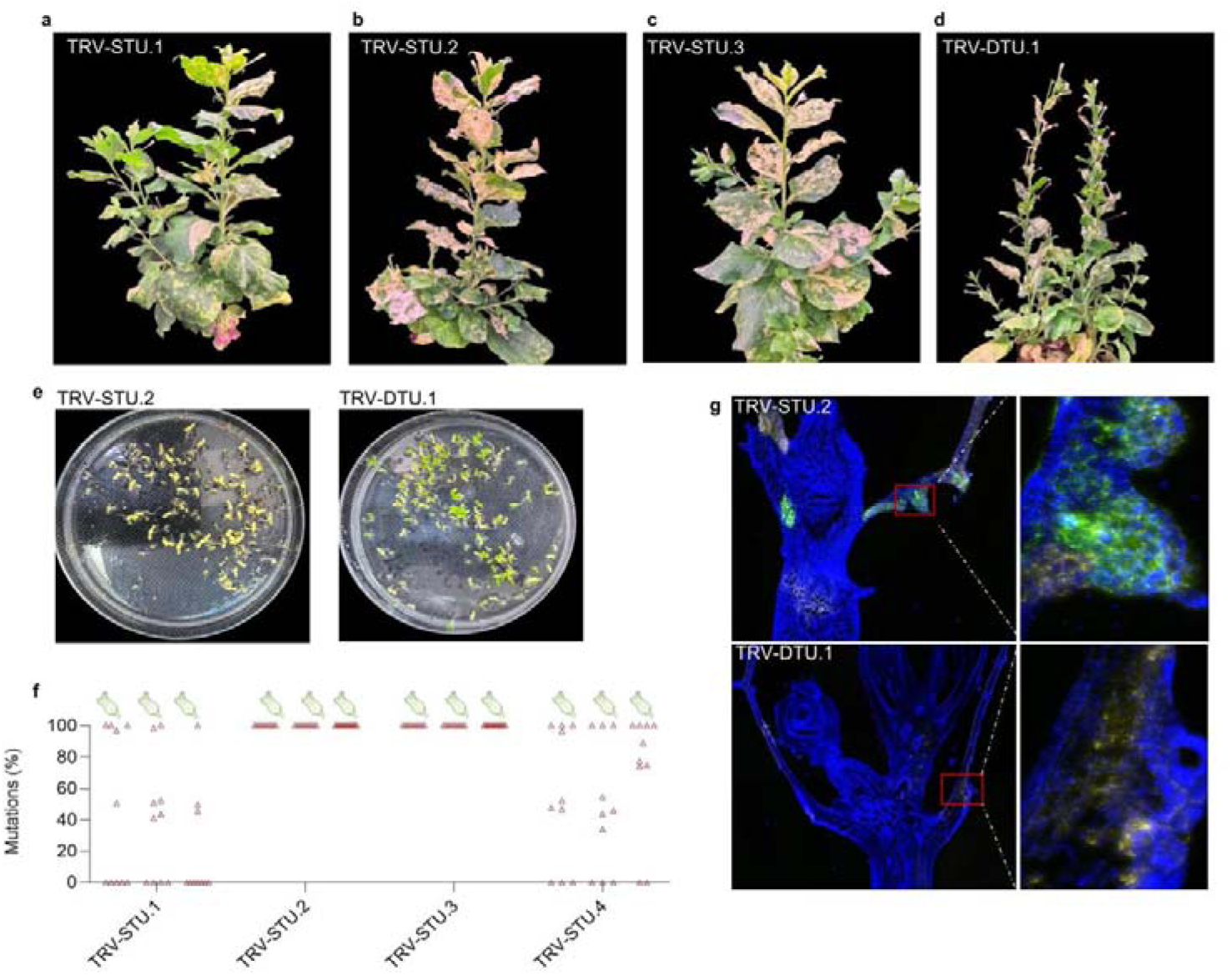
STU architecture enables VIGE-mediated heritable genome editing. **a-d**, Phenotypes of *N. benthamiana* plants approximately 2 months after infiltration with the four configurations. **e,** Phenotypes of progeny derived from chlorotic capsules collected from plants infected with TRV-STU.2 and TRV-DTU.1. **f,** Genome editing frequencies in seedlings derived from chlorotic capsules of plants infected with the four TnpB configurations. Ten seedlings per capsule were randomly selected for amplicon deep sequencing, and at least three independent capsules were analyzed for each construct. **g,** RNA FISH of floral tissues collected from plants infected with TRV-STU.2 and TRV-DTU.1. The two tissues were sectioned from the same paraffin-embedded sample. Green fluorescence indicates *ISDra2 TnpB* RNA signals, whereas yellow fluorescence indicates CP RNA signals.

### The STU architecture promotes retention of TnpB editing cargo during systemic viral propagation

To investigate the basis underlying the difference in heritable genome editing efficiency between TRV-STU.2 and TRV-DTU.1, we performed RNA fluorescence in situ hybridization (FISH) using probes targeting the CP and TnpB sequences (Supplementary Table 1). Sixty days after infiltration, floral tissues derived from the primary shoot apices of flowering plants were collected and analyzed (Supplementary Fig. 2). In TRV-STU.2-infected plants, both CP and TnpB signals were readily detected in developing leaf primordia, indicating the sustained presence of the viral vector-derived transcripts and retention of the inserted TnpB sequence during long-term systemic propagation (Fig. 3g). In contrast, in TRV-DTU.1-infected plants, only CP signals were detected, whereas no detectable TnpB signal was observed (Fig. 3g). These results suggest that although the TRV-DTU.1 construct can induce visible editing-associated phenotypes, the inserted TnpB region may undergo progressive instability or depletion during long-term viral propagation, resulting in reduced accumulation of TnpB transcripts in systemic tissues. Interestingly, RNA FISH analysis did not detect TRV signals in the shoot apical meristem (Fig. 3g), consistent with our observation that newly emerged tissues derived from the apical meristem initially lacked chlorotic phenotypes. In contrast, chlorotic phenotypes were frequently observed in stem tissues and at the axillary regions located at the base of older leaf primordia, where axillary meristems are established. Axillary meristems retain the developmental capacity of the SAM and can generate new leaf primordia, lateral shoots, and reproductive structures, providing a potential route for the generation of heritable genome-edited progeny.

### Combining VIGS and VIGE enables phenotype-guided screening of heritable genome edits

Although VIGE has enabled efficient genome editing in plants, most applications have focused on genes with readily observable phenotypes, such as *ChlH* or *PDS*. During our experiments, we found that even *ChlH*-targeted plants frequently produced both chlorotic and non-chlorotic capsules, and progeny derived from non-chlorotic capsules rarely carried detectable edits. These observations revealed a major challenge in identifying heritable edited progeny, particularly for target genes lacking visible phenotypes, which often requires extensive genotyping. To overcome this limitation, we sought to establish a visual screening workflow that enables efficient identification of heritable genome-edited progeny (Fig. 4a-b). We first introduced the RUBY reporter cassette into the TRV2 vector backbone (Fig. 4a). RUBY expression provided a rapid visual indicator of successful viral delivery and gene expression in infiltrated leaves (Fig. 4b, c). Previous studies have shown that TRV vectors are predominantly used for virus-induced gene silencing (VIGS), largely due to their simplicity: insertion of a 200–300 bp fragment of an endogenous gene into the viral genome is sufficient to induce gene silencing (Senthil-Kumar and Mysore, 2011). In the present study, the compact nature of our STU architecture—being shorter than previously reported dual-expression TnpB and reRNA constructs—provides additional cargo capacity for the incorporation of functional modules (Nagalakshmi et al., 2026). We therefore explored whether TRV could be engineered to simultaneously support TnpB-mediated genome editing and silencing of the endogenous *NbPDS* gene, whose knockdown produces a readily scorable albino phenotype that serves as a visual selection marker. Notably, *NbPDS* silencing is rarely inherited into the next generation, making it suitable as a transient screening indicator. To validate this strategy, we constructed three dual-function vectors by introducing *NbPDS* silencing fragments into the TRV-STU.2, TRV-STU.3, and TRV-DTU.1 backbones, generating VS-STU.2, VS-STU.3, and VS-DTU.1, respectively (Fig. 4a). To compare the performance of different STU configurations in the VIGE system, we selected *NbChlH* as the target gene and evaluated genome editing efficiencies in infiltrated leave (Fig. 4c). The three vectors exhibited distinct editing profiles. Notably, incorporation of the *NbPDS* silencing fragment generally reduced editing efficiency, with VS-DTU.1 showing the most pronounced decrease, potentially due to its increased transcript length compared with VS-STU.2 and VS-STU.3 (Fig. 4d). At 14 days post infiltration (dpi), VS-STU.2 and VS-STU.3 induced clear albino phenotypes in systemic tissues (Fig. 4c), whereas VS-DTU.1 produced weak or undetectable albino phenotypes. Therefore, subsequent analyses focused primarily on VS-STU.2 and VS-STU.3.

**Fig. 4.**
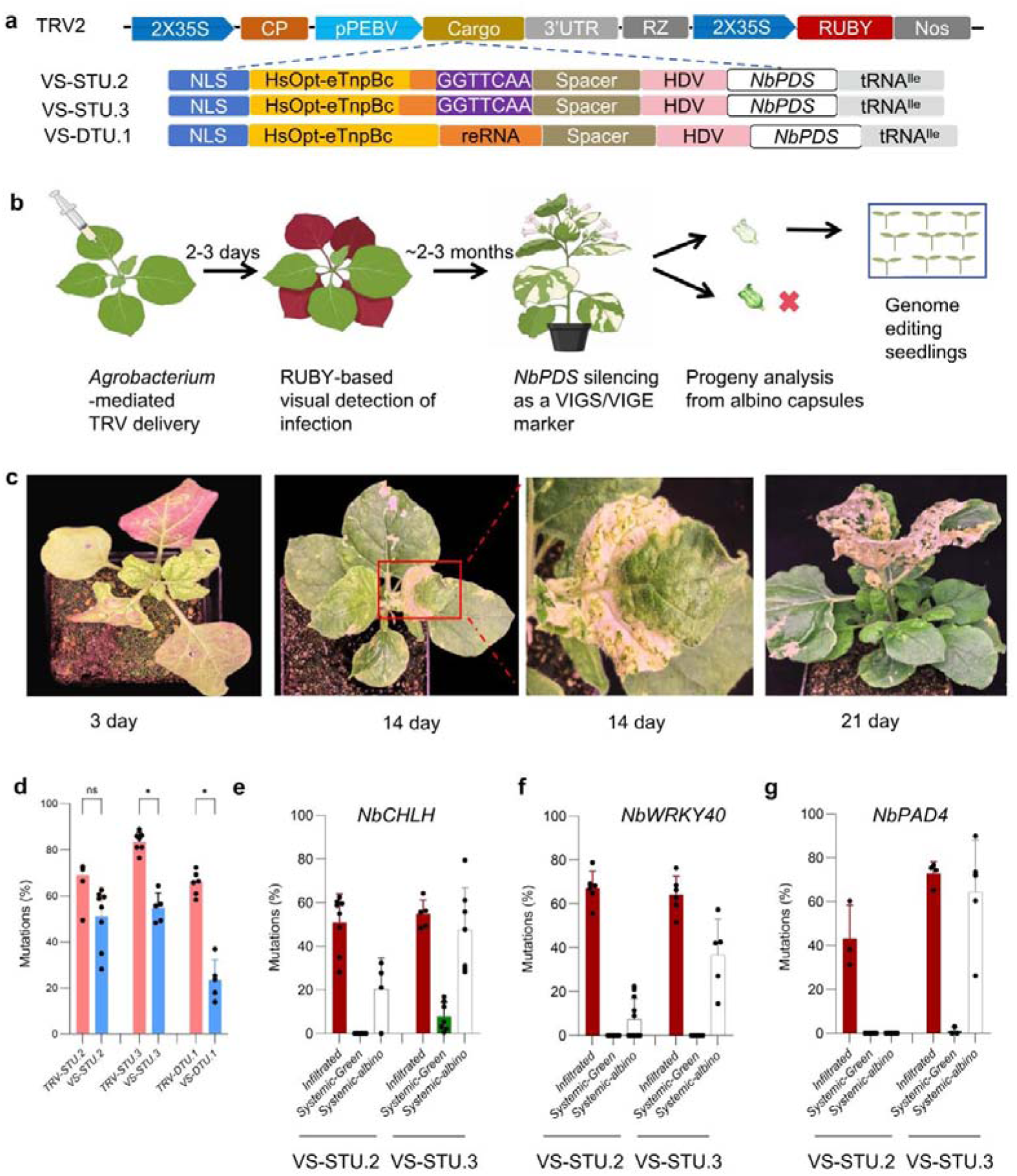
Establishment of a visual screening system for VIGE through VIGS. **a,** Schematic representation of TRV2-derived vector systems carrying STU-based TnpB expression configurations and an *NbPDS*-derived fragment for VIGE. **b,** Workflow for visual screening of genome-edited progeny. **c,** Phenotypic progression of *N. benthamiana* plants infected with VS-STU.3. Images were taken at 3, 14 and 21 days after TRV infiltration. RUBY expression was observed in infiltrated leaves at 3 days after infiltration, whereas *NbPDS* silencing-associated albino phenotypes became visible at approximately 14 days after infiltration. **d,** Comparison of genome editing efficiencies mediated by VS-STU.2, VS-STU.3 and VS-DTU.1 in infiltrated leaves. Data were analysed using an unpaired t-test with a two-tailed P value. ns, P > 0.05; *P < 0.05; Each point represents an independent biological replicate (n > 3). **e–g,** Genome editing efficiencies of three endogenous target genes mediated by VS-STU.2 and VS-STU.3 in infiltrated leaves, systemic leaves, and systemic leaves lacking visible albino phenotypes.

To further evaluate the robustness and applicability of the system beyond genes with visible phenotypes, we selected two endogenous *N. benthamiana* genes, *NbWRKY40* and *NbPAD4*, whose disruption does not cause obvious phenotypic changes under normal growth conditions. Corresponding STU-based constructs were generated and introduced into *N. benthamiana* plants via *Agrobacterium*-mediated infiltration. Infiltrated leaves were collected at 3 dpi, whereas systemic tissues were harvested at 21 dpi, including both green and albino leaves, for genome editing analysis. Strikingly, VS-STU.2 and VS-STU.3 displayed distinct editing patterns in systemic tissues. In infiltrated leaves, both systems exhibited consistently high genome editing efficiencies across three target sites, indicating that the two configurations were efficiently delivered and functional in infiltrated leaves (Fig. 4e-g). However, their performance diverged markedly in systemic tissues collected at 21 dpi (Fig. 4e-g). Although albino systemic leaves were observed in both cases, all tested targets in the VS-STU.2 system showed minimal editing activity, whereas VS-STU.3 maintained high editing efficiencies comparable to those detected in infiltrated leaves (Fig. 4e-g). Notably, within the VS-STU.3 system, systemic leaves exhibiting albino phenotypes displayed substantially higher genome editing efficiencies than those without visible albino phenotypes, indicating that virus-induced genome silencing (VIGS) and VIGE mediated by the VS-STU.3 system are closely associated and occur in a coordinated manner (Fig. 3e-g). These results suggest that differences in codon optimization may substantially influence viral mobility and systemic stability. To investigate the underlying basis, we predicted the RNA secondary structures of the corresponding transcripts. Although VS-STU.2 and VS-STU.3 differed by only 108 nucleotides in sequence composition, their predicted RNA secondary structures exhibited pronounced differences (Supplementary Fig. 3). Notably, for VS-STU.2, the inconsistent predictions between the MFE and centroid structures suggest that the transcript may adopt a dynamic ensemble of conformations rather than a single dominant structure, which may contribute to its distinct functional behavior and reduced genome editing activity in systemic tissues. Furthermore, exchanging the positions of the *NbPDS*-derived fragment and tRNA^ile^ within the VS-STU.3 configuration resulted in a significant reduction in genome editing efficiency in systemic leaves (Supplementary Fig. 4). Previous studies have also shown that distinct codon optimization strategies can substantially affect genome editing performance (Nagamatsu et al., 2007). Together, these findings suggest that viral performance in VIGE systems is not determined solely by cargo size. Instead, RNA sequence composition and structural features may represent additional, previously underappreciated determinants regulating viral stability, mobility, and systemic editing activity.

Next, we evaluated whether visible phenotypic selection could facilitate the identification of heritable genome-edited progeny. Green and albino capsules were separately collected from either independent plants or the same individual plants (Fig. 5a–d), and progeny derived from these capsules were subjected to genome editing analysis (Fig. 5e-g). Progeny from green capsules showed no detectable editing events across all tested loci (Fig. 5g). By contrast, albino capsules efficiently enriched for genome-edited progeny. At the *NbChlH* locus, 80% of analyzed progeny seedlings carried detectable genome edits, and 26.7% exhibited chlorotic phenotypes, indicating that the visible albino phenotype was associated with successful heritable editing events (Fig. 5g). Similarly, for the non-phenotypic target genes *NbWRKY40* and *NbPAD4*, 100% and 73.3% of progeny seedlings derived from albino capsules carried detectable genome edits, respectively. Notably, 40% of *NbWRKY40*-edited progeny exhibited tetra-allelic mutations (Fig. 3g–h), further demonstrating efficient enrichment of edited individuals through the albino capsule-based selection strategy. Together, these results demonstrate that the combined VIGS–VIGE strategy enables efficient identification and enrichment of genome-edited progeny, including targets lacking intrinsic visible phenotypes, thereby providing a practical solution for downstream screening in plant genome editing applications.

**Fig. 5.**
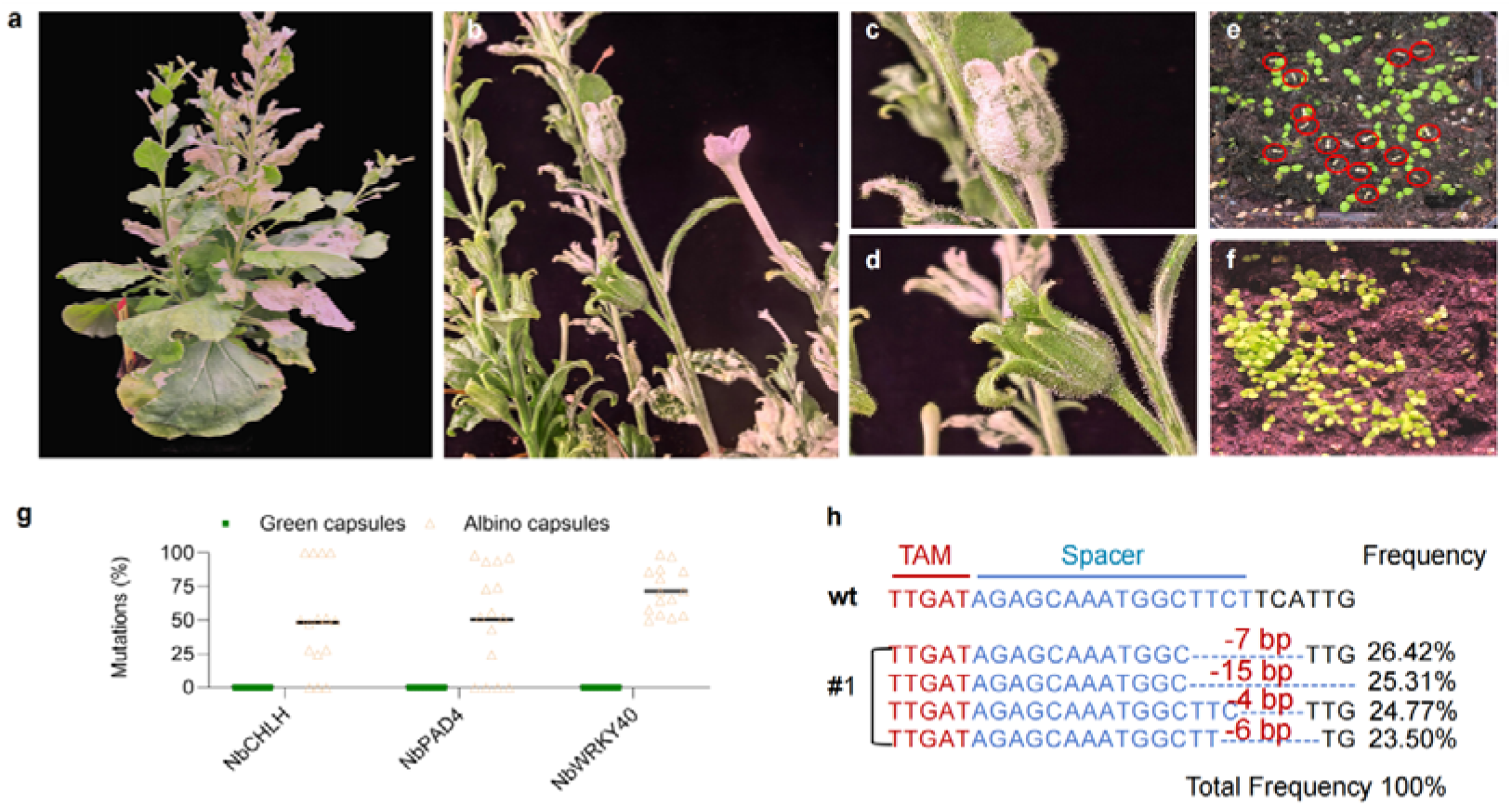
VIGS-based screening enriches VIGE-derived heritable genome-edited progeny. **a-b,** Phenotypes of *N. benthamiana* plants two months after infiltration with VS-STU.3 targeting *NbChlH*. A subset of seed capsules displayed albino-associated phenotypes. **c-d,** Representative capsules with albino (**c**) or non-albino (**d**) phenotypes from VS-STU.3-infected plants targeting *NbChlH.* **e-f,** Phenotypes of progeny seedlings derived from albino seed capsules (**e**) and non-albino seed capsules (**f**), red circles indicate seedlings displaying chlorotic phenotypes. **g,** Comparison of genome editing frequencies between progeny seedlings derived from albino and non-albino seed capsules. At least 15 progeny seedlings were randomly selected from each seed capsule for amplicon deep sequencing analysis. **h,** Genotypes of tetra-allelic *NbWRKY40* mutants generated by the VIGS-VIGE strategy.

### No detectable off-targets and viral sequences in STU-edited progeny

Moreover, to evaluate potential off-target effects of the STU system, we analyzed three target sites in rice and three target sites in *N. benthamiana* exhibiting relatively high editing efficiencies. No detectable mutations were detected at the predicted off-target loci, suggesting a high degree of target specificity for the STU-mediated genome editing system (Supplementary Table 2). RT–PCR analysis showed that no TRV2 coat protein (CP) transcripts were detected in any progeny seedlings, indicating that viral transcripts were undetectable in edited progeny seedlings (Supplementary Fig. 5).

## Discussion

Although biochemical studies have demonstrated that TnpB can process its own transcript to generate reRNA molecules (Nety et al., 2023), this native overlapping configuration has not yet been shown to support efficient genome editing in vivo (Luo et al., 2025). Consequently, previous TnpB-based genome editing systems have generally adopted separate expression strategies for the TnpB nuclease and reRNA components (Altae-Tran et al., 2023; Karvelis et al., 2021; Li et al., 2024; Lv et al., 2024; Molla et al., 2024; Wang et al., 2024), as the functional potential of the naturally overlapping reRNA–coding architecture in eukaryotic systems remained unclear. Here, we demonstrate that this compact organization can be functionally maintained in plant cells, highlighting the potential of native single-transcript architectures as an alternative design strategy for genome editing. Whether this architecture can be similarly harnessed in mammalian cells remains to be determined. If transferable, the STU configuration could provide substantial advantages for delivery platforms with stringent cargo limitations, such as adeno-associated virus (AAV)-based genome editing, by minimizing the size of the editing cassette and increasing the capacity for additional regulatory or functional elements.

Previous studies have shown that directly linking full-length eTnpBc with target-specific reRNA, combined with additional optimizations such as nuclear localization signals, can achieve high editing efficiencies in plants (Nagalakshmi et al., 2026). Our results demonstrate that the STU architecture can achieve comparable genome editing outcomes without requiring such extensive optimization, suggesting that transcript compactness may provide advantages for VIGE applications by reducing cargo size and potentially improving viral delivery and/or systemic movement.

Previous studies on VIGE optimization have primarily focused on minimizing the size of genome editing cargos to overcome the limited packaging capacity of viral vectors (Mikhaylova, 2025; Qiao et al., 2026; Steinberger and Voytas, 2025). More recent studies have further demonstrated that codon optimization can substantially enhance TnpB-mediated VIGE efficiency (Nagalakshmi et al., 2026). Our findings extend these observations by showing that viral genome engineering performance is influenced by additional RNA-level determinants beyond cargo size and protein expression. Although VS-STU.2 and VS-STU.3 differed by only 108 nucleotides, they exhibited markedly distinct editing outcomes in systemic tissues (Fig. 4), suggesting that subtle changes in RNA sequence composition can substantially influence viral behavior (Fig. 4 and Supplementary Fig. 3). The predicted differences in RNA secondary structures, together with the reduced editing efficiency caused by repositioning the *NbPDS*-derived fragment within the VS-STU.3 configuration (Supplementary Fig. 3), indicate that RNA structural features may affect transcript stability, viral replication, systemic movement, or accessibility of the embedded editing components. These RNA-level determinants have received limited attention in previous VIGE studies but may represent an important regulatory layer for optimizing viral genome editing platforms.

## Methods

### Vector Construction

The vectors used for rice genome editing were constructed based on the pCXUN backbone as previously described (Sun et al., 2016). The codon-optimized *ISDra2 TnpB* gene and reRNA scaffold sequences were synthesized by Sangon Biotech (Shanghai, China). Target-specific reRNA sequences were introduced by overlapping PCR. For the STU vectors, the overlapping TnpB-reRNA cassette was driven by the maize Ubiquitin promoter. For the DTU vectors, codon-optimized *ISDra2 TnpB* and reRNA were independently expressed under the control of the maize *Ubiquitin* promoter and the rice OsU3 promoter, respectively. All DNA fragments were assembled into the linearized pCXUN backbone using Seamless Cloning (TransGen Biotech).

The TRV1 and TRV2 vectors were constructed based on previously described systems (Ellison et al., 2020). DNA fragments used for TRV vector construction, including different optimized versions of *ISDra2 TnpB*, were synthesized by Sangon Biotech (Shanghai, China). The corresponding fragments were cloned into the AatII/XmaI-digested TRV2 backbone using Seamless Cloning (TransGen Biotech). Target-specific sequences were introduced into TRV-based editing constructs by overlapping PCR. Primers used for vector construction are listed in Supplementary Table 3. The Information of target sites used in this study are listed in Supplementary Table 4. All generated plasmids were verified by Sanger sequencing of the inserted regions.

### Rice stable transformation and soybean hairy root transformation

For rice transformation, all constructs were introduced into *Oryza sativa* cv. Zhonghua11 using *Agrobacterium tumefaciens* strain EHA105-mediated transformation performed by EDGENE Biotechnology (Wuhan) Co., Ltd. (Wuhan, Hubei, China).

For soybean hairy root transformation, transgenic hairy roots were generated following a previously described protocol with modifications (Cao et al., 2025). Briefly, seven-day-old soybean seedlings were wounded at the hypocotyl region and inoculated with *Agrobacterium rhizogenes* strain K599. Inoculated seedlings were transferred to sterile vermiculite and maintained under high-humidity conditions. Hairy root formation was observed approximately two weeks after inoculation.

### Plant growth conditions and *Agrobacterium*-mediated TRV delivery

*N. benthamiana* plants were grown in a controlled growth chamber at 26 °C under a 12 h light/12 h dark cycle, with a light intensity of 100 μE m^−2^ s^−1^ and 50% relative humidity. TRV1 and TRV2-derived vectors were introduced into *Agrobacterium tumefaciens* strains GV3101(P19) and GV3101, respectively. Transformants were selected on LB agar medium supplemented with kanamycin (50 μg ml^−1^) and rifampicin (25 μg ml^−1^) and incubated at 28 °C for 2 days. Five colonies randomly selected for each construct, streaked onto fresh LB agar plates containing the same antibiotics, and incubated overnight at 28 °C. Positive transformants were confirmed by colony PCR and stored as glycerol stocks at −80 °C until further use.

For agroinfiltration, glycerol stocks were recovered from −80 °C and streaked onto LB agar plates containing kanamycin (50 μg ml^−1^) and rifampicin (25 μg ml^−1^), followed by overnight incubation at 28 °C. *Agrobacterial* cells were collected from the plates and resuspended in agroinfiltration medium containing 10 mM MgCl₂, 10 mM 2-(N-morpholine)-ethanesulfonic acid (MES), and 250 μM 3′,5′-dimethoxy-4′-hydroxyacetophenone (acetosyringone). The bacterial suspension was adjusted to an OD_600_ of 0.6 and incubated with gentle shaking for at least 3 h at room temperature prior to infiltration. Equal volumes of TRV1 and TRV2-derived bacterial suspensions were mixed at a 1:1 ratio and infiltrated into three diagonally opposite leaves of 2.5–3-week-old *N. benthamiana* plants using a 1 ml needleless syringe. Following infiltration, plants were maintained under laboratory growth conditions at 24 °C under a 12 h light/12 h dark photoperiod, with a light intensity of 100 μE m^−2^ s^−1^ and 50% relative humidity.

### Genome editing efficiency analysis

Genome editing efficiencies in transgenic rice and TRV-infected plants were determined by amplicon deep sequencing. Genomic DNA was extracted from transgenic rice tissues, TRV-infiltrated leaves, and systemic leaves using the same DNA extraction method. For TRV-mediated genome editing analysis, leaf discs approximately 8-10 mm in diameter were collected from the infiltrated regions at 3 dpi or from upper systemic leaves at 3-5 weeks dpi prior to DNA extraction. Amplicon sequencing was performed following a previously described protocol (Sun et al., 2024). Briefly, PCR products encompassing the target regions were prepared and subjected to high-throughput sequencing using the Hi-TOM platform at the State Key Laboratory of Rice Biology and Breeding, China National Rice Research Institute, Chinese Academy of Agricultural Sciences (Hangzhou, China).

### RNA structure analysis

Predicted secondary structures of processed RNA scaffolds were generated using the RNAfold web server from the ViennaRNA package (http://rna.tbi.univie.ac.at/). Minimum free energy (MFE) and centroid structures were calculated using default parameters.

### Nanopore full-length transcript sequencing and analysis

Full-length transcript sequencing was performed using the Oxford Nanopore long-read sequencing platform. Raw sequencing reads were subjected to quality control to remove low-quality reads and short fragments, and the resulting clean reads were used for downstream analysis. A reference sequence containing the complete exogenous expression cassette was used for read alignment. Clean reads were mapped to the reference sequence using minimap2, and uniquely mapped reads with a mapping quality score ≥20 were retained for subsequent analysis. To specifically identify transgene-derived transcripts and reduce nonspecific background signals, reads containing a conserved 3′ terminal sequence motif (ATCACGCGACTTTAGTCGTG for rice and TTTAGTCGTGTGAGGTTCAA for *N. benthamiana*) were extracted and defined as target transcript reads. These reads were subsequently remapped to the reference sequence, and the alignment start position (POS) of each read was extracted from the resulting BAM files to evaluate 5′-end integrity. Reads with POS=1 were considered full-length transcripts containing the complete 5′ region, whereas reads with POS >1 were classified as 5′-truncated transcripts, with the deletion length calculated as POS-1.The distribution and relative abundance of 5′ truncation sites were determined based on the frequency of reads initiating at each genomic position. For visualization and association with functional regions, truncation sites were further grouped into 20-bp intervals to generate 5’-end deletion profiles.

### RNA fluorescence in situ hybridization

RNA fluorescence in situ hybridization (RNA FISH) targeting TRV *coat protein* (*CP*) and *ISDra2 TnpB* transcripts was performed by Suzhou Dynamic Biosystems Co., Ltd. (Suzhou, China). Sections were hybridized with specific fluorescent probes targeting *CP* and *ISDra2 TnpB* transcripts. After probe hybridization and washing, fluorescence signals were visualized using a Leica DM6B microscope (Leica, Germany). Cell walls were counterstained with Calcofluor White (CFW) and imaged simultaneously. Images showing merged signals of *CP, ISDra2 TnpB*, and CFW staining were generated, and individual channels for each probe and CFW staining were also presented.

### RT–PCR

Total RNA was extracted from TRV-infiltrated leaves and progeny seedlings using TRIzol reagent (TransGen Biotech) and treated with RNase-free DNase I (TransGen Biotech) to eliminate genomic DNA contamination. First-strand cDNA was synthesized using SuperScript III reverse transcriptase (TransGen Biotech) according to the manufacturer’s instructions. PCR amplification was subsequently performed using the synthesized cDNA as the template with primers specifically designed for the TRV CP gene and the endogenous *N. benthamiana PP2A* gene.

### Off-target analysis

Potential off-target sites for the STU-mediated genome editing system were predicted using the online tool (http://skl.scau.edu.cn/) described previously (Xie et al., 2017), based on the presence of a ‘TTGAT’ TAM and four to six nucleotide mismatches relative to the target sequences of three high-editing-efficiency loci (*OsYSA*, *OsWaxy*, and *OsHMBPP*). The genomic regions surrounding the predicted off-target sites were amplified from 15 independent seedlings and analyzed using the same amplicon deep sequencing approach as described for target site analysis.

## Data analysis

The data were analyzed using GraphPad Prism 8.0 software, and the figures were further processed using Adobe Photoshop and Adobe Illustrator software.

## Authors contributions

Y. S and D.L conceived the study and designed the experiments. R.W., H.L., S.J., S.B., and X.C performed the experiments. Y. S., D.L., R.W., H.L. and S.J., analysed the data. Y.S and D.L wrote and revised the manuscript. All authors read and approved the final manuscript. All authors read and approved the manuscript.

## Acknowledgements

This work was funded by the Major Science and Technology Project of Inner Mongolia Autonomous Region (Grant No. 2026ZD0081 to D.L), National Natural Science Foundation of China (Grant No. 32370431, 32160111 to Y.S; 32160143 to D.L), The Inner Mongolia Natural Science Foundation (Grant No. 2025MS03007 to Y.S).

## Conflict of interest

The authors declare no conflict of interest.

## Notes

### Competing Interest Statement

The authors have declared no competing interest.

### Summary of Updates

Added funding information and revised the introduction section.

