## Supplementary Figures and tables for "A native single-transcript TnpB architecture enables efficient virus-induced genome editing and visual screening of heritable progeny harboring phenotypically silent edits"

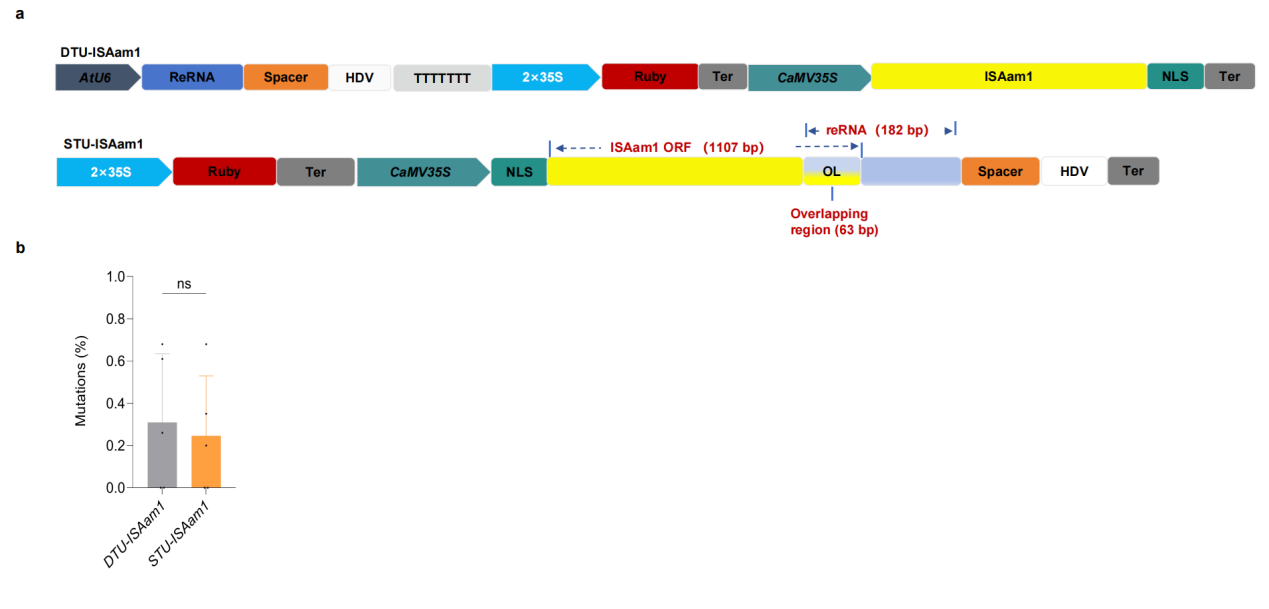

**Supplementary Fig. 1 A native single-transcript ISAam1 TnpB system enables genome editing in soybean hairy roots.**

**a**, Schematic representation of STU and DTU constructs based on ISAam1 TnpB used for genome editing in soybean hairy roots. In the DTU-ISAam1 construct, ISAam1 TnpB and reRNA were independently expressed under the control of the CaMV 35S and *Arabidopsis* U6 promoters, respectively. In the STU-ISAam1 construct, ISAam TnpB and reRNA were incorporated into a single transcriptional unit driven by the CaMV 35S promoter. **b,** Comparison of genome editing efficiencies between STU-ISAam1 and DTU-ISAam1 constructs targeting the soybean endogenous gene *GmFAD2-1A*.

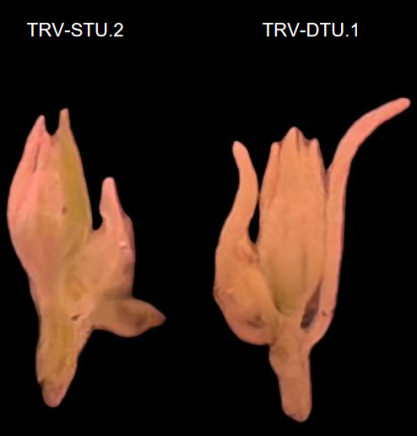

**Supplementary Fig. 2 Floral tissues isolated from flowering plants 60 days after infiltration.**

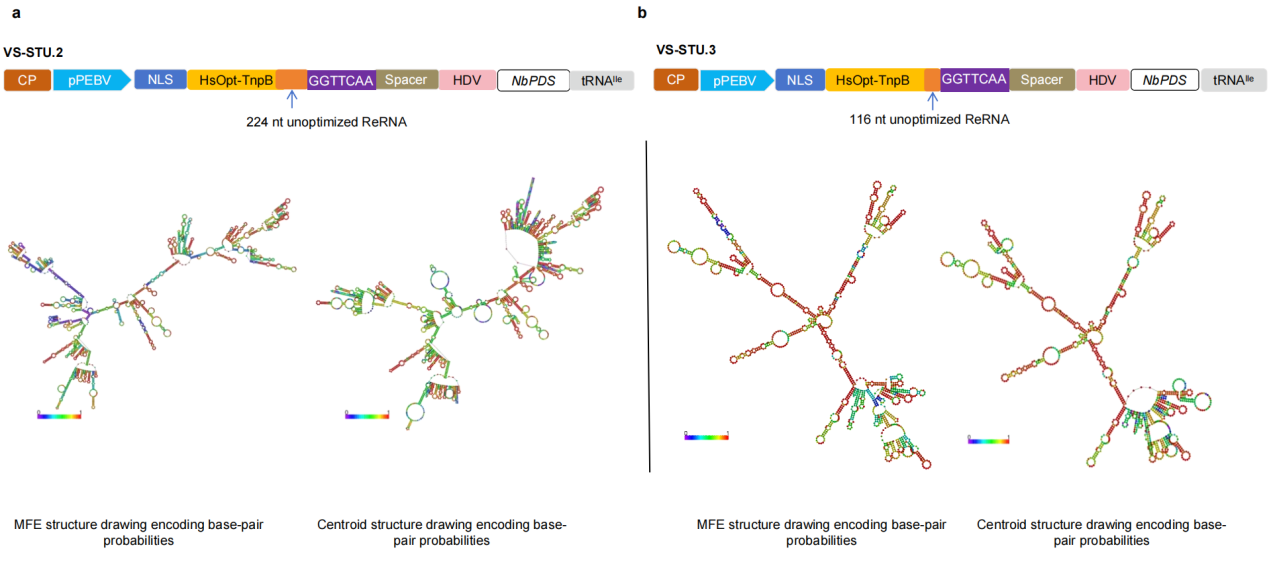

**Supplementary Fig. 3 Predicted RNA secondary structures of VS-STU.2 and VS-STU.3 transcripts.**

a-b**,** Predicted minimum free energy (MFE) and centroid secondary structures of the VS-STU.2 (**a**) and VS-STU.3 (**b**) transcripts.

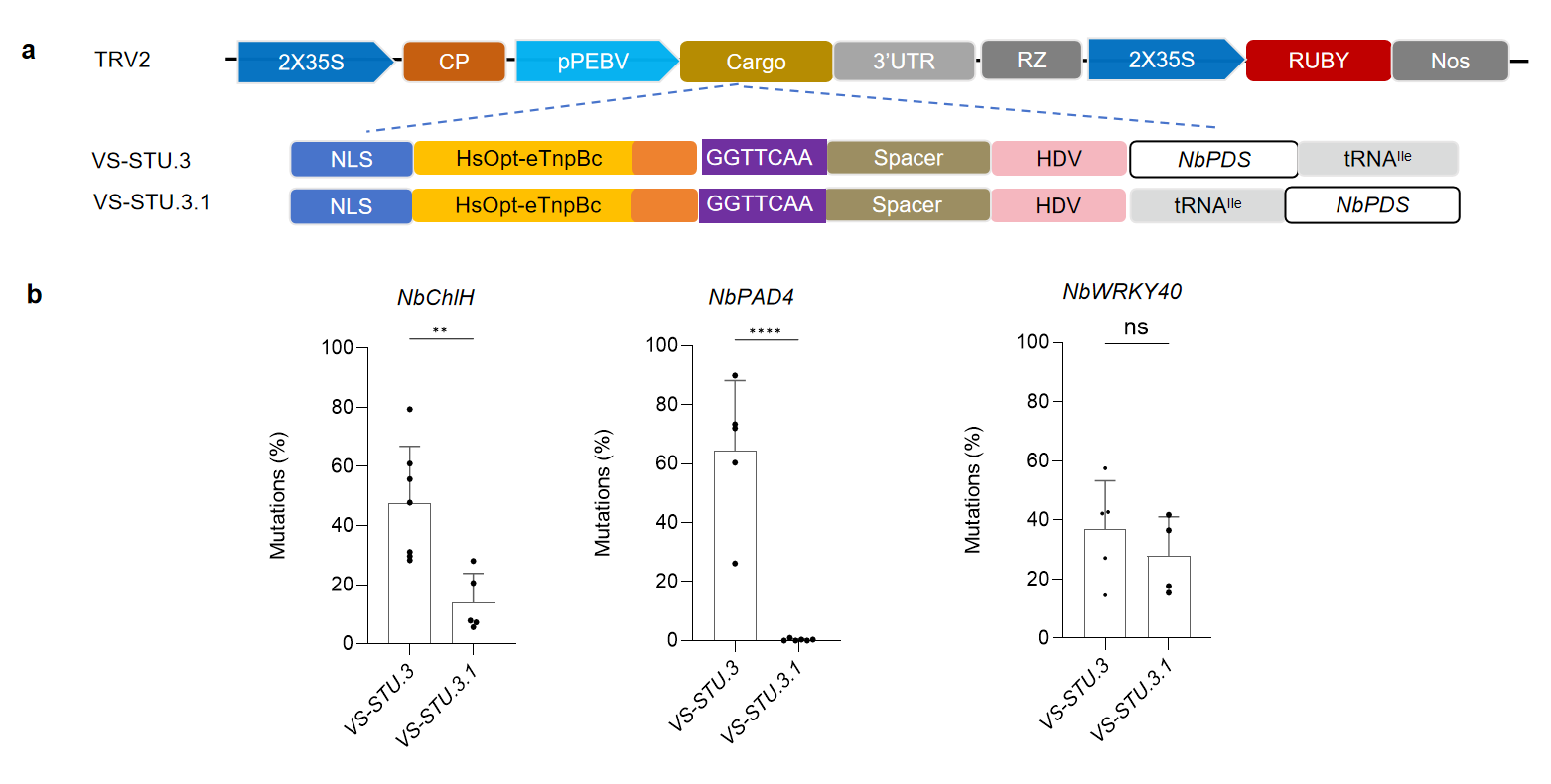

**Supplementary Fig. 4 Effect of RNA element organization on genome editing efficiency in systemic leaves of *N. benthamiana*.**

**a,** Schematic representation of TRV2-derived vector systems carrying STU-based TnpB expression configurations and an NbPDS-derived fragment for virus-induced gene silencing. For the VS-STU.3.1 the NbPDS-derived fragment and tRNA^Ile^ were exchanged compared with VS-STU.3. **b,** Comparison of genome editing efficiencies between VS-STU.3 and VS-STU.3.1 constructs targeting the targeting three genes. Data were analysed using an unpaired t-test with a two-tailed Pvalue. ns, P > 0.05; *P < 0.05; Each point represents a biological replicate from an independent experiment (n > 3).

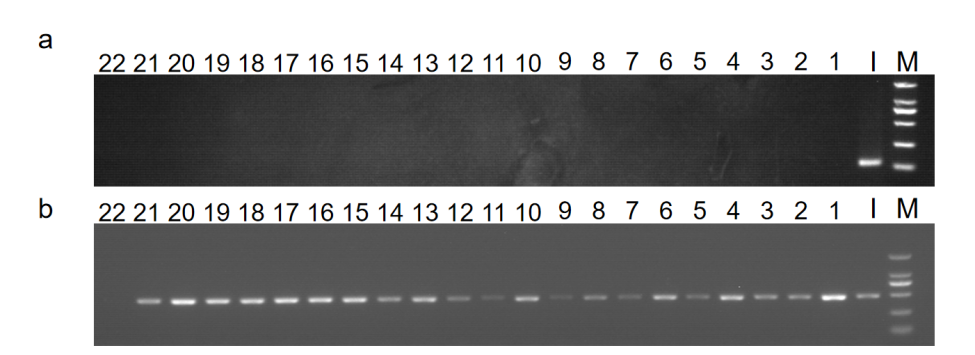

**Supplementary Fig. 5 Absence of detectable TRV-derived sequences in progeny seedlings of edited *N. benthamiana* plants.**

**a**, RT–PCR analysis showing the absence of detectable TRV2 coat protein (CP) transcripts in progeny seedlings derived from edited plants. **b**, RT–PCR amplification of the endogenous PP2A gene as an internal control. M, 2 kb DNA ladder; I, infiltrated leaf samples; lanes 1–22, individual progeny seedlings; lane 23, water control.

**Supplementary Tables**

**Supplementary Table 1. Sequences of RNA FISH probes used in this study.**

| **Target genes** | **Probe Pairs** | **Target sequences (5'-3')** |
| --- | --- | --- |
| CP | 5 | GGTCCTGCTGACTTGATGGACGATTCTTGGGT |
|  |  | GCACTACAGTCTGGTAGAGATGAGATCACTGG |
|  |  | GACTCACGGGCTAACAGTGCTCTTGGTGTGAT |
|  |  | AACACAACGGTTACGACGAACCAAGGGAGTAC |
|  |  | GGTGGAGCAGCTGCTAGTTCATCTGCACCGCC |
| ISDra2 TnpB | 8 | GCACAAACTGAGCTGATAAACCGAACACTGGG |
|  |  | GAATCGCTGCGTACAAGGAAAGCGGTAAGGGT |
|  |  | CTCCTGGCTGTCAGAAGTGGATAAATTCGCGT |
|  |  | CAGATAGGAGAGGGGAGGCTTAAACTGCCGAA |
|  |  | CTGTGCGAGGTCGAAATCCCTTACCTCCCAGC |
|  |  | CTATCGGAGTACGCTTAAGCGAATTCGGAAGG |
|  |  | AAGACTAAGCTGGCAAGGATTCACAAGCGCAT |
|  |  | CCGACAATATGCGCAAGAATAGAAGGCTGGCG |

**Supplemental Table 2. Analysis of potential off-target effects**

| **Target** | **Off-target site** | **The putative off-target site (5'-3')** | **No. of individual plants sequenced** | **Gene** | **No. of plants with mutations** |
| --- | --- | --- | --- | --- | --- |
| *OsYAS* | OFF-T1 | TTGATTGACAATaTTTGCTCACTTG | 15 |  | 0 |
|  | OFF-T2 | TTGATTGACAAgCTTTGaTCtCTTG | 15 | OsZH11G0715392700.01 | 0 |
| *OsWaxy* | OFF-T1 | cTGAT TACAAAaACAACtAaAgGCG | 15 | OsZH11G0180000290.01 | 0 |
|  | OFF-T2 | TTGATcACAAAttCAAaACAGATGCa | 15 |  | 0 |
| *OsHMBPP* | OFF-T1 | TTaATCTCAAACACCTTCTGcAgCa | 15 | OsZH11G0305602600.01 | 0 |
|  | OFF-T2 | TTtcTCcCAAACACCTTCTGaAACG | 15 |  | 0 |
| *NbChlH* | OFF-T1 | TTGgTGAAGAGATCTCgtAgCA | 15 | Niben101Scf02046 | 0 |
|  | OFF-T2 | TTGATGAAGAGATaTCTGAAaA | 15 | Niben101Scf02547 | 0 |
|  | OFF-T3 | TTGAgGAAaAtAaCTCTGAACA | 15 | Niben101Scf05203 | 0 |
| *NbPAD4* | OFF-T1 | TTGATCACTAGAgTTATaACA | 15 | Niben101Scf05467 | 0 |
|  | OFF-T2 | TTGATCACTtGtTTTATCACA | 15 | Niben101Scf10150 | 0 |
|  | OFF-T3 | TTGATaACTAtATTTATCAaA | 15 | Niben101Scf00077 | 0 |
| *NbWRKY40* | OFF-T1 | TTGATAGAaCAAATGGCTTCT | 15 | Niben101Ctg16115 | 0 |
|  | OFF-T2 | TTGATAGAGCAcATGGgTTtT | 15 | Niben101Scf00461 | 0 |
|  | OFF-T3 | TTGATAGAGaAAATacCTTCT | 15 | Niben101Scf00567 | 0 |

Note: the TAM motif is underlined; mismatched bases are indicated in lowercase letters.

**Supplementary Table 3. The oligonucleotide primers used in this study**

| **Name** | **Sequence 5'-3'** | **Description** |
| --- | --- | --- |
| Hi-OsPDS-T1-F1 | GAGTACGGTGTGCATGGAAGGAACACTCCATGA | Sequencing primer for *OsPDS-T1* site |
| Hi-OsPDS-T1-R1 | GGATGCTGGATGGACCACCAATACGATGTAC |  |
| Hi-OsPDS-T2-F1 | GAGTACGGTGTGCGGCTACTCATTTTAACTGAC | Sequencing primer for *OsPDS-T2* site |
| Hi-OsPDS-T2-R1 | GGATGCTGGATGGGGTCATATGTGTTCTTCAG |  |
| Hi-OsWaxy-F1 | GAGTACGGTGTGCGTTTGGGGAAAGACCGGTG | Sequencing primer for *OsWaxy* site |
| Hi-OsWaxy-R1 | GGATGCTGGATGGAGCATCAATACACGAATTAG |  |
| Hi-OsHMBPP-F1 | GAGTACGGTGTGCCAGAAGGCCCTATCACAATC | Sequencing primer for *OsHMBPP* site |
| Hi-OsHMBPP-R1 | GGATGCTGGATGGTCGCTTGTGCACTGGTTCTA |  |
| Hi-OsCKX2-F1 | GAGTACGGTGTGCTCGGTGATCTCTCTGCAAAG | Sequencing primer for *OsCKX2* site |
| Hi-OsCKX2-R1 | GGATGCTGGATGGTCGTGGACGCGGTCCAGGAA |  |
| Hi-OsCERK1-F1 | GAGTACGGTGTGCTCTTGTAATAGGTTATGAAC | Sequencing primer for *OsHMBPP* site |
| Hi-CERK1-R1 | GGATGCTGGATGGAAAAGGAACACGTGAAATCG |  |
| Hi-OsPi21-F1 | GAGTACGGTGTGCCGCAGAGGAGTACTGCATCG | Sequencing primer for *OsPi21* site |
| Hi-OsPi21-R1 | GGATGCTGGATGGGTCCTCCGGAGGCTTCTCG |  |
| Hi-OsYSA-F1 | GAGTACGGTGTGCTAACGCGCTCTGCAAAGTAG | Sequencing primer for *OsYSA* site |
| Hi-OsYSA-R1 | GGATGCTGGATGGATATGTAATTGTACTTCGAG |  |
| Hi-NbPAD4-F1 | GAGTACGGTGTGCATCCTTGCAATTGATTTATC | Sequencing primer for *NbPAD4* site |
| Hi-NbPAD4-F2 | GAGTACGGTGTGCACTCGCAACCTTAATAGTGC |  |
| Hi-NbPAD4-R1 | GGATGCTGGATGGTAGCAAGTCTGATCCTGTTC |  |
| Hi-NbPAD4-R2 | GGATGCTGGATGGCACTTGTCAATCACAGAATC |  |
| Hi-NbPAD4-R3 | GGATGCTGGATGGGAAAATCATAAGGCAGTTGA |  |
| Hi-NbWRKY40-F1 | GAGTACGGTGTGCACAACAGAAGGTGATCACT | Sequencing primer for *NbWRKY40* site |
| Hi-NbWRKY40-F2 | GAGTACGGTGTGCAAGGTGGCAATAGCCACAGG |  |
| Hi-NbWRKY40-R1 | GGATGCTGGATGGGTTTACCATCTACTTGTCTG |  |
| Hi-NbWRKY40-R2 | GGATGCTGGATGGGGAAATTGACTGGTGTGCAG |  |
| Hi-NbWRKY40-R3 | GGATGCTGGATGGTGCTAGCCTATTCATAAATG |  |
| Hi-NbChlH-F1 | GAGTACGGTGTGCGTTAGTCTTGAAGAGCTTG | Sequencing primer for *NbChlH* site |
| Hi-NbChlH-F2 | GAGTACGGTGTGCGGGTTAACCAGGTGGAACCT |  |
| Hi-NbChlH-F3 | GAGTACGGTGTGCTGTTACAGACACGTTAGGAC |  |
| Hi-NbChlH-R1 | GGATGCTGGATGGTGTTGAAAATAAAACAGTAAG |  |
| Hi-NbChlH-R2 | GGATGCTGGATGGCGAGCTCAGCAACCATCTTC |  |
| Hi-NbChlH-R3 | GGATGCTGGATGGAGTTTTGGTCTTCTGGCTCG |  |
| TRV2-Nb-F | GGGAATTTCCTTTACCATTGACGTCAGTGTCGTTGG | Forward and reverse primers used for TRV vector construction, paired with target gene-specific primers for amplification. |
| TRV2-Nb-R | AATGTCTTCGGGACATGCCCGGGCCTCGAGTGCTTCCGGCGGGGCTCGAACCCGCGACC |  |
| TRV-NbChlH-F | GAAGAGATCTCTGAACACCCGGCCGGCATGGTCCCAGCCT | Construction of the *NbChlH*-targeting vector for genome editing |
| TRV-NbChlH-R | GGGTGTTCAGAGATCTCTTCTTGAACCTCACACGACTAAA |  |
| TRV-NbPAD4-F | CACTAGATTTATCACAGGCCGGCATGGTCCCAGCCT | Construction of the *NbPAD4-*targeting vector for genome editing |
| TRV-NbPAD4-R | GGCCTGTGATAAATCTAGTGTTGAACCTCACACGACTAAA |  |
| TRV-NbWRKY40-F | AGAGCAAATGGCTTCTGGCCGGCATGGTCCCAGCCT | Construction of the *NbWRKY40*-targeting vector for genome editing |
| TRV-NbWRKY40-R | GGCCAGAAGCCATTTGCTCTTTGAACCTCACACGACTAAA |  |
| OsU3-reRNA-F | TGTCGTTTCCCGCCTTCAGTTTGTAATTCATCCAGGTCTCCAAG | Forward and reverse primers used for construction of the reRNA expression cassette in DTU vectors, which were paired with target gene-specific forward and reverse primers, respectively. |
| OsU3-reRNA-R | ACACTGATAGTTTAAAAAAAGTCCCATTCGCCATGCCGAAGCATGTTGCCCAGCCGGCG |  |
| Rice DTU-F | TGGTGTTACTTCTGCAGCCCGGGGGATCCCCAATACTATGGCCCCAAAGAAGAAGC | Forward and reverse primers used to integrate TnpB into the vector backbone for construction of rice DTU vectors. |
| Roce DTU-R | TTGAACGATCGGGGAAATTCGGATCCCCAATACTAAAAAAAGTCCCATTCGCCATGCCG |  |
| OsYAS-F | TGACAATCTTTGCTCACTTGGGCCGGCATGGTCCCAGCCT | Construction of the OsYAS-targeting vector for genome editing |
| OsYAS-R | CAAGTGAGCAAAGATTGTCATTGAACCTCACACGACTAAA |  |
| OsCERK1-F | GTCCCGATGTATGTATACTGGGCCGGCATGGTCCCAGCCT | Construction of the *OsCERK1*-targeting vector for genome editing |
| OsCERK1-R | CAGTATACATACATCGGGACTTGAACCTCACACGACTAAA |  |
| OsPDS-T1-F | TTTCCTGAAACATTGCCTGCGGCCGGCATGGTCCCAGCCT | Construction of the *OsPDS-T1*-targeting vector for genome editing |
| OsPDS-T1-R | GCAGGCAATGTTTCAGGAAATTGAACCTCACACGACTAAA |  |
| OsPDS-T2-F | ATCTTGAAGCTTCTTGTACCGGCCGGCATGGTCCCAGCCT | Construction of the *OsPDS-T2*-targeting vector for genome editing |
| OsPDS-T2-R | GGTACAAGAAGCTTCAAGATTTGAACCTCACACGACTAAA |  |
| OsHMBPP-F | CTCAAACACCTTCTGTAACGGGCCGGCATGGTCCCAGCCT | Construction of the *OsHMBPP-*targeting vector for genome editing |
| OsHMBPP-R | CGTTACAGAAGGTGTTTGAGTTGAACCTCACACGACTAAA |  |
| OsPi21-F | GATCTTGCCGGCCTTGCACCGGCCGGCATGGTCCCAGCCT | Construction of the *OsPi21*-targeting vector for genome editing |
| OsPi21-R | GGTGCAAGGCCGGCAAGATCTTGAACCTCACACGACTAAA |  |
| OsCKX2-F | TTTATTTGCAGAGGATGGATGGCCGGCATGGTCCCAGCCT | Construction of the *OsCKX2*-targeting vector for genome editing |
| OsCKX2-R | ATCCATCCTCTGCAAATAAATTGAACCTCACACGACTAAA |  |
| OsWaxy-F | TACAAAGACAACCAGATGCGGGCCGGCATGGTCCCAGCCT | Construction of the *OsWaxy*-targeting vector for genome editing |
| OsWaxy-R | CGCATCTGGTTGTCTTTGTATTGAACCTCACACGACTAAA |  |

Note:for the same target, multiple primer pairs are often available, and forward and reverse primers may be alternated to prevent cross-contamination, which could lead to inaccurate experimental data. Additionally, certain primers may be shared across different target sites. Sequencing can only proceed if no amplification bands are observed in the negative control for each experiment.

**Supplementary Table 4. Information of target sites used in this study.**

| **Species** | **TAM** | **Target** | **Sequence (5'-3')** |
| --- | --- | --- | --- |
| Rice | TTGAT | *OsPDS-T1* | TTTCCTGAAACATTGCCTGC |
|  | TTGAT | *OsPDS-T2* | ATCTTGAAGCTTCTTGTACC |
|  | TTGAT | *OsHMBPP* | CTCAAACACCTTCTGTAACG |
|  | TTGAT | *OsYSA* | TGACAATCTTTGCTCACTTG |
|  | TTGAT | *OsWAXY* | TACAAAGACAACCAGATGCG |
|  | TTGAT | *OsCKX2* | TTTATTTGCAGAGGATGGAT |
|  | TTGAT | *OsCERK1* | GTCCCGATGTATGTATACTG |
|  | TTGAT | *OsPi21* | GATCTTGCCGGCCTTGCACC |
| Soybean | TTTAA | *GmFAD2-1A* | GTACTTGGAAAACCATGCAA |
| *Nicotiana benthamiana* | TTGAT | *NbChlH* | AGAACGTGCTGGGCAAGT |
|  | TTGAT | *NbPAD4* | GGGCGTGGCGCCATGGCT |
|  | TTGAT | *NbWRKY40* | TAATATTGGATCCACCGA |

**Supplementary sequence 1**

**Cassette of STU for rice genome editing**

Ubi-NLS-TnpB (reRNA-Stop Codon (TGA))- GGTTCAA-Spacer-HDV-Nos

CTGCAGTGCAGCGTGACCCGGTCGTGCCCCTCTCTAGAGATAATGAGCATTGCATGTCTAAGTTATAAAAAATTACCACATATTTTTTTTGTCACACTTGTTTGAAGTGCAGTTTATCTATCTTTATACATATATTTAAACTTTACTCTACGAATAATATAATCTATAGTACTACAATAATATCAGTGTTTTAGAGAATCATATAAATGAACAGTTAGACATGGTCTAAAGGACAATTGAGTATTTTGACAACAGGACTCTACAGTTTTATCTTTTTAGTGTGCATGTGTTCTCCTTTTTTTTTGCAAATAGCTTCACCTATATAATACTTCATCCATTTTATTAGTACATCCATTTAGGGTTTAGGGTTAATGGTTTTTATAGACTAATTTTTTTAGTACATCTATTTTATTCTATTTTAGCCTCTAAATTAAGAAAACTAAAACTCTATTTTAGTTTTTTTATTTAATAATTTAGATATAAAATAGAATAAAATAAAGTGACTAAAAATTAAACAAATACCCTTTAAGAAATTAAAAAAACTAAGGAAACATTTTTCTTGTTTCGAGTAGATAATGCCAGCCTGTTAAACGCCGTCGACGAGTCTAACGGACACCAACCAGCGAACCAGCAGCGTCGCGTCGGGCCAAGCGAAGCAGACGGCACGGCATCTCTGTCGCTGCCTCTGGACCCCTCTCGAGAGTTCCGCTCCACCGTTGGACTTGCTCCGCTGTCGGCATCCAGAAATGCGTGGCGGAGCGGCAGACGTGAGCCGGCACGGCAGGCGGCCTCCTCCTCCTCTCACGGCACGGCAGCTACGGGGGATTCCTTTCCCACCGCTCCTTCGCTTTCCCTTCCTCGCCCGCCGTAATAAATAGACACCCCCTCCACACCCTCTTTCCCCAACCTCGTGTTGTTCGGAGCGCACACACACACAACCAGATCTCCCCCAAATCCACCCGTCGGCACCTCCGCTTCAAGGTACGCCGCTCGTCCTCCCCCCCCCCCCCTCTCTACCTTCTCTAGATCGGCGTTCCGGTCCATGGTTAGGGCCCGGTAGTTCTACTTCTGTTCATGTTTGTGTTAGATCCGTGTTTGTGTTAGATCCGTGCTGCTAGCGTTCGTACACGGATGCGACCTGTACGTCAGACACGTTCTGATTGCTAACTTGCCAGTGTTTCTCTTTGGGGAATCCTGGGATGGCTCTAGCCGTTCCGCAGACGGGATCGATTTCATGATTTTTTTTGTTTCGTTGCATAGGGTTTGGTTTGCCCTTTTCCTTTATTTCAATATATGCCGTGCACTTGTTTGTCGGGTCATCTTTTCATGCTTTTTTTTGTCTTGGTTGTGATGATGTGGTCTGGTTGGGCGGTCGTTCTAGATCGGAGTAGAATTCTGTTTCAAACTACCTGGTGGATTTATTAATTTTGGATCTGTATGTGTGTGCCATACATATTCATAGTTACGAATTGAAGATGATGGATGGAAATATCGATCTAGGATAGGTATACATGTTGATGCGGGTTTTACTGATGCATATACAGAGATGCTTTTTGTTCGCTTGGTTGTGATGATGTGGTGTGGTTGGGCGGTCGTTCATTCGTTCTAGATCGGAGTAGAATACTGTTTCAAACTACCTGGTGTATTTATTAATTTTGGAACTGTATGTGTGTGTCATACATCTTCATAGTTACGAGTTTAAGATGGATGGAAATATCGATCTAGGATAGGTATACATGTTGATGTGGGTTTTACTGATGCATATACATGATGGCATATGCAGCATCTATTCATATGCTCTAACCTTGAGTACCTATCTATTATAATAAACAAGTATGTTTTATAATTATTTTGATCTTGATATACTTGGATGATGGCATATGCAGCAGCTATATGTGGATTTTTTTAGCCCTGCCTTCATACGCTATTTATTTGCTTGGTACTGTTTCTTTTGTCGATGCTCACCCTGTTGTTTGGTGTTACTTCTGCAGCCCGGGGGATCCCCAATACTATGGCCCCAAAGAAGAAGCGCAAGGTCATCCGCtAcAAGGCCTTCGTGGTGAGGCTCTACCCGAACGCCGCCCAGACCGAGCTCATTAATAGGACCCTCGGCTCTGCCAGGTTCGTCTACAATCATTTCCTCGCGAGAAGGATTGCCGCTTACAAGGAGTCAGGCAAGGGCCTGACATACGGCCAGACATCTTCAGAGCTCACACTCCTTAAGCAGGCTGAGGAGACATCTTGGCTGTCTGAGGTGGATAAGTTCGCCCTCCAGAATAGCTTGAAGAATCTGGAAACCGCTTACAAAAATTTCTTCAGGACAGTGAAGCAGTCTGGCAAGAAGGTTGGCTTCCCTAGGTTCaagAAGAAGCGCACAGGCGAGTCATATAGGACTCAATTCACAAATAACAATATTCAGATCGGCGAGGGCAGGCTCAAGCTCCCGAAGCTCGGCTGGGTCAAGACCAAGGGCCAGCAAGACATCCAGGGCAAGATCCTGAACGTGACCGTGCGCCGGATCCACGAGGGCCACTACGAGGCCAGCGTGCTCTGCGAGGTGGAGATCCCGTACCTGCCGGCCGCCCCGAAGTTCGCGGCGGGCGTGGACcTCGGCATCAAGGACTTCGCCATCGTGACCGATGGCGTGCGCTTCAAGCACGAGCAGAACCCGAAGTACTACCGCAGCACCCTCAAGCGCaTCCGCAAGGCGCAGCAGACCCTCTCCCGCCGCAAGAAGGGCAGCGCCCGCTACGGCAAGGCCAAGACCAAGCTCGCCCGGATCCACAAGAGGATCGTGAACAAGCGCCAGGATTTCCTCCACAAGCTCACCACCTCCCTCGTGCGCGAGTACGAGATCATCGGCACCGAGCATCTGAAGCCGGATAACATGAGGAAGAACCGCAGGCTCGCCCTGAGCATCAGCGATGCCGGCTGGGGCGAGTTCATCCGCCAGCTCGAGTACAAGGCCGCGTGGTACGGCAGGCTCGTGAGCAAGGTGTCCCCGTACTTCCCAAGCTCCCAGCTCTGCCACGACTGCGGATTCAAGAATCCCGAAGTGAAGAATCTTGCCGTCCGTACATGGACTTGCCCGAACTGTGGGGAAACCCATGACCGAGACGAGAACGCTGCGCTGAACATTCGGCGTGAAGCGTTGGTGGCTGCGGGAATCTCAGACACCTTAAACGCTCATGGAGGCTATGTCAGACCTGCTTCGGCGGGCAATGGTCTGCGAAGTGAGAATCACGCGACTTTAGTCGTGTGAGGTTCAATATCCAAGACCAGAGCTAGAggccggcatggtcccagcctcctcgctggcgccggctgggcaacatgcttcggcatggcgaatgggacTTTTTTTAGTATTGGGGATCCGAATTTCCCCGATCGTTCAAACATTTGGCAATAAAGTTTCTTAAGATTGAATCCTGTTGCCGGTCTTGCGATGATTATCATATAATTTCTGTTGAATTACGTTAAGCATGTAATAATTAACATGTAATGCATGACGTTATTTATGAGATGGGTTTTTATGATTAGAGTCCCGCAATTATACATTTAATACGCGATAGAAAACAAAATATAGCGCGCAAACTAGGATAAATTATCGCGCGCGGTGTCATCTATGTTACTAGATC

**Supplementary sequence 2**

**DTU cassette for TnpB expression in rice genome editing**

Ubi-NLS-Tnpb -Nos

CTGCAGTGCAGCGTGACCCGGTCGTGCCCCTCTCTAGAGATAATGAGCATTGCATGTCTAAGTTATAAAAAATTACCACATATTTTTTTTGTCACACTTGTTTGAAGTGCAGTTTATCTATCTTTATACATATATTTAAACTTTACTCTACGAATAATATAATCTATAGTACTACAATAATATCAGTGTTTTAGAGAATCATATAAATGAACAGTTAGACATGGTCTAAAGGACAATTGAGTATTTTGACAACAGGACTCTACAGTTTTATCTTTTTAGTGTGCATGTGTTCTCCTTTTTTTTTGCAAATAGCTTCACCTATATAATACTTCATCCATTTTATTAGTACATCCATTTAGGGTTTAGGGTTAATGGTTTTTATAGACTAATTTTTTTAGTACATCTATTTTATTCTATTTTAGCCTCTAAATTAAGAAAACTAAAACTCTATTTTAGTTTTTTTATTTAATAATTTAGATATAAAATAGAATAAAATAAAGTGACTAAAAATTAAACAAATACCCTTTAAGAAATTAAAAAAACTAAGGAAACATTTTTCTTGTTTCGAGTAGATAATGCCAGCCTGTTAAACGCCGTCGACGAGTCTAACGGACACCAACCAGCGAACCAGCAGCGTCGCGTCGGGCCAAGCGAAGCAGACGGCACGGCATCTCTGTCGCTGCCTCTGGACCCCTCTCGAGAGTTCCGCTCCACCGTTGGACTTGCTCCGCTGTCGGCATCCAGAAATGCGTGGCGGAGCGGCAGACGTGAGCCGGCACGGCAGGCGGCCTCCTCCTCCTCTCACGGCACGGCAGCTACGGGGGATTCCTTTCCCACCGCTCCTTCGCTTTCCCTTCCTCGCCCGCCGTAATAAATAGACACCCCCTCCACACCCTCTTTCCCCAACCTCGTGTTGTTCGGAGCGCACACACACACAACCAGATCTCCCCCAAATCCACCCGTCGGCACCTCCGCTTCAAGGTACGCCGCTCGTCCTCCCCCCCCCCCCCTCTCTACCTTCTCTAGATCGGCGTTCCGGTCCATGGTTAGGGCCCGGTAGTTCTACTTCTGTTCATGTTTGTGTTAGATCCGTGTTTGTGTTAGATCCGTGCTGCTAGCGTTCGTACACGGATGCGACCTGTACGTCAGACACGTTCTGATTGCTAACTTGCCAGTGTTTCTCTTTGGGGAATCCTGGGATGGCTCTAGCCGTTCCGCAGACGGGATCGATTTCATGATTTTTTTTGTTTCGTTGCATAGGGTTTGGTTTGCCCTTTTCCTTTATTTCAATATATGCCGTGCACTTGTTTGTCGGGTCATCTTTTCATGCTTTTTTTTGTCTTGGTTGTGATGATGTGGTCTGGTTGGGCGGTCGTTCTAGATCGGAGTAGAATTCTGTTTCAAACTACCTGGTGGATTTATTAATTTTGGATCTGTATGTGTGTGCCATACATATTCATAGTTACGAATTGAAGATGATGGATGGAAATATCGATCTAGGATAGGTATACATGTTGATGCGGGTTTTACTGATGCATATACAGAGATGCTTTTTGTTCGCTTGGTTGTGATGATGTGGTGTGGTTGGGCGGTCGTTCATTCGTTCTAGATCGGAGTAGAATACTGTTTCAAACTACCTGGTGTATTTATTAATTTTGGAACTGTATGTGTGTGTCATACATCTTCATAGTTACGAGTTTAAGATGGATGGAAATATCGATCTAGGATAGGTATACATGTTGATGTGGGTTTTACTGATGCATATACATGATGGCATATGCAGCATCTATTCATATGCTCTAACCTTGAGTACCTATCTATTATAATAAACAAGTATGTTTTATAATTATTTTGATCTTGATATACTTGGATGATGGCATATGCAGCAGCTATATGTGGATTTTTTTAGCCCTGCCTTCATACGCTATTTATTTGCTTGGTACTGTTTCTTTTGTCGATGCTCACCCTGTTGTTTGGTGTTACTTCTGCAGCCCGGGGGATCCCCAATACTATGGCCCCAAAGAAGAAGCGCAAGGTCATCCGCtAcAAGGCCTTCGTGGTGAGGCTCTACCCGAACGCCGCCCAGACCGAGCTCATTAATAGGACCCTCGGCTCTGCCAGGTTCGTCTACAATCATTTCCTCGCGAGAAGGATTGCCGCTTACAAGGAGTCAGGCAAGGGCCTGACATACGGCCAGACATCTTCAGAGCTCACACTCCTTAAGCAGGCTGAGGAGACATCTTGGCTGTCTGAGGTGGATAAGTTCGCCCTCCAGAATAGCTTGAAGAATCTGGAAACCGCTTACAAAAATTTCTTCAGGACAGTGAAGCAGTCTGGCAAGAAGGTTGGCTTCCCTAGGTTCaagAAGAAGCGCACAGGCGAGTCATATAGGACTCAATTCACAAATAACAATATTCAGATCGGCGAGGGCAGGCTCAAGCTCCCGAAGCTCGGCTGGGTCAAGACCAAGGGCCAGCAAGACATCCAGGGCAAGATCCTGAACGTGACCGTGCGCCGGATCCACGAGGGCCACTACGAGGCCAGCGTGCTCTGCGAGGTGGAGATCCCGTACCTGCCGGCCGCCCCGAAGTTCGCGGCGGGCGTGGACcTCGGCATCAAGGACTTCGCCATCGTGACCGATGGCGTGCGCTTCAAGCACGAGCAGAACCCGAAGTACTACCGCAGCACCCTCAAGCGCaTCCGCAAGGCGCAGCAGACCCTCTCCCGCCGCAAGAAGGGCAGCGCCCGCTACGGCAAGGCCAAGACCAAGCTCGCCCGGATCCACAAGAGGATCGTGAACAAGCGCCAGGATTTCCTCCACAAGCTCACCACCTCCCTCGTGCGCGAGTACGAGATCATCGGCACCGAGCATCTGAAGCCGGATAACATGAGGAAGAACCGCAGGCTCGCCCTGAGCATCAGCGATGCCGGCTGGGGCGAGTTCATCCGCCAGCTCGAGTACAAGGCCGCGTGGTACGGCAGGCTCGTGAGCAAGGTGTCCCCGTACTTCCCAAGCTCCCAGCTCTGCCACGACTGCGGATTCAAGAATCCCGAAGTGAAGAATCTTGCCGTCCGTACATGGACTTGCCCGAACTGTGGGGAAACCCATGACCGAGACGAGAACGCTGCGCTGAACATTCGGCGTGAAGCGTTGGTGGCTGCGGGAATCTCAGACACCTTAAACGCTCATGGAGGCTATGTCAGACCTGCTTCGGCGGGCAATGGTCTGCGAAGTGAGAATCACGCGACTTTAGTCGTGTGAAGTATTGGGGATCCGAATTTCCCCGATCGTTCAAACATTTGGCAATAAAGTTTCTTAAGATTGAATCCTGTTGCCGGTCTTGCGATGATTATCATATAATTTCTGTTGAATTACGTTAAGCATGTAATAATTAACATGTAATGCATGACGTTATTTATGAGATGGGTTTTTATGATTAGAGTCCCGCAATTATACATTTAATACGCGATAGAAAACAAAATATAGCGCGCAAACTAGGATAAATTATCGCGCGCGGTGTCATCTATGTTACTAGATC

**Supplementary sequence 3**

**DTU cassette for reRNA expression in rice genome editing**

**OsU3-reRNA-Spacer-HDV-Poly T**

gtaattcatccaggtctccaagttctaggattttcagaactgcaacttattttatcaaggaatctttaaacatacgaacagatcacttaaagttcttctgaagcaacttaaagttatcaggcatgcatggatcttggaggaatcagatgtgcagtcagggaccatagcacaagacaggcgtcttctactggtgctaccagcaaatgctggaagccgggaacactgggtacgttggaaaccacgtgatgtgaagaagtaagataaactgtaggagaaaagcatttcgtagtgggccatgaagcctttcaggacatgtattgcagtatgggccggcccattacgcaattggacgacaacaaagactagtattagtaccacctcggctatccacatagatcaaagctgatttaaaagagttgtgcagatgatccgtggcaGATTCAAGAATCCCGAAGTGAAGAATCTTGCCGTCCGTACATGGACTTGCCCGAACTGTGGGGAAACCCATGACCGAGACGAGAACGCTGCGCTGAACATTCGGCGTGAAGCGTTGGTGGCTGCGGGAATCTCAGACACCTTAAACGCTCATGGAGGCTATGTCAGACCTGCTTCGGCGGGCAATGGTCTGCGAAGTGAGAATCACGCGACTTTAGTCGTGTGAGGTTCAACAGTCTTCTCAGAAGACGTggccggcatggtcccagcctcctcgctggcgccggctgggcaacatgcttcggcatggcgaatgggacTTTTTT

**Supplementary sequence 4**

**TRV-STU.2 sequence for Nicotiana benthamiana genome editing**

NLS-STU.2 TnpB (reRNA (224 bp unoptimized)-Stop Codon (TGA)- GGTTCAA-Spacer)-HDV-tRNA^ile^

ATGGCCCCAAAGAAGAAGCGCAAGGTCATACGCtatAAGGCATTTGTGGTTCGGCTCTACCCGAATGCGGCACAAACTGAGCTGATAAACCGAACACTGGGGTCTGCACGATTTGTATACAATCATTTCCTTGCTCGAAGAATCGCTGCGTACAAGGAAAGCGGTAAGGGTCTTACGTACGGGCAAACTTCTAGTGAGCTTACTCTGCTGAAGCAAGCAGAGGAAACCTCCTGGCTGTCAGAAGTGGATAAATTCGCGTTGCAGAATAGCTTGAAAAACCTTGAAACGGCCTATAAAAACTTCTTTAGAACAGTTAAGCAGTCAGGAAAAAAGGTAGGTTTCCCTCGGTTCaaaAAGAAGCGCACGGGGGAGTCATACCGCACGCAGTTTACGAACAACAACATTCAGATAGGAGAGGGGAGGCTTAAACTGCCGAAACTGGGGTGGGTCAAAACTAAGGGTCAGCAGGATATCCAAGGCAAAATATTGAATGTTACTGTACGCCGCATCCATGAAGGCCATTATGAAGCTTCAGTGCTGTGCGAGGTCGAAATCCCTTACCTCCCAGCTGCTCCGAAATTCGCGGCAGGAGTAGACCTGGGGATTAAAGACTTTGCGATCGTCACCGATGGTGTGCGATTTAAGCATGAGCAAAATCCTAAATACTATCGGAGTACGCTTAAGCGAATTCGGAAGGCCCAACAGACTCTGTCCAGGAGAAAGAAAGGAAGCGCTAGATATGGCAAGGCTAAGACTAAGCTGGCAAGGATTCACAAGCGCATCGTGAATAAGAGACAAGATTTTCTGCACAAACTGACCACTTCCCTTGTGCGCGAGTATGAAATAATCGGGACTGAACACCTGAAGCCCGACAATATGCGCAAGAATAGAAGGCTGGCGCTCAGCATCTCAGATGCTGGATGGGGTGAATTCATAAGGCAGTTGGAGTACAAAGCTGCTTGGTATGGGAGACTCGTAAGTAAAGTTTCCCCATATTTTCCTTCTAGTCAACTCTGTCACGACTGCGGATTCAAGAATCCCGAAGTGAAGAATCTTGCCGTCCGTACATGGACTTGCCCGAACTGTGGGGAAACCCATGACCGAGACGAGAACGCTGCGCTGAACATTCGGCGTGAAGCGTTGGTGGCTGCGGGAATCTCAGACACCTTAAACGCTCATGGAGGCTATGTCAGACCTGCTTCGGCGGGCAATGGTCTGCGAAGTGAGAATCACGCGACTTTAGTCGTGTGAGGTTCAAGAAGAGATCTCTGAACACCCggccggcatggtcccagcctcctcgctggcgccggctgggcaacatgcttcggcatggcgaatgggacccgtagctcagttggttagagcgttggtcttatgagccgaaggtcgcgggttcgagccccgccggaagca

**Supplementary sequence 5**

**TRV-STU.3 sequence for Nicotiana benthamiana genome editing**

NLS-STU.3 TnpB (reRNA (116 bp unoptimized)-Stop Codon (TGA)- GGTTCAA-Spacer)-HDV-tRNA^ile^

ATGGCCCCAAAGAAGAAGCGCAAGGTCATACGCtatAAGGCATTTGTGGTTCGGCTCTACCCGAATGCGGCACAAACTGAGCTGATAAACCGAACACTGGGGTCTGCACGATTTGTATACAATCATTTCCTTGCTCGAAGAATCGCTGCGTACAAGGAAAGCGGTAAGGGTCTTACGTACGGGCAAACTTCTAGTGAGCTTACTCTGCTGAAGCAAGCAGAGGAAACCTCCTGGCTGTCAGAAGTGGATAAATTCGCGTTGCAGAATAGCTTGAAAAACCTTGAAACGGCCTATAAAAACTTCTTTAGAACAGTTAAGCAGTCAGGAAAAAAGGTAGGTTTCCCTCGGTTCaaaAAGAAGCGCACGGGGGAGTCATACCGCACGCAGTTTACGAACAACAACATTCAGATAGGAGAGGGGAGGCTTAAACTGCCGAAACTGGGGTGGGTCAAAACTAAGGGTCAGCAGGATATCCAAGGCAAAATATTGAATGTTACTGTACGCCGCATCCATGAAGGCCATTATGAAGCTTCAGTGCTGTGCGAGGTCGAAATCCCTTACCTCCCAGCTGCTCCGAAATTCGCGGCAGGAGTAGACCTGGGGATTAAAGACTTTGCGATCGTCACCGATGGTGTGCGATTTAAGCATGAGCAAAATCCTAAATACTATCGGAGTACGCTTAAGCGAATTCGGAAGGCCCAACAGACTCTGTCCAGGAGAAAGAAAGGAAGCGCTAGATATGGCAAGGCTAAGACTAAGCTGGCAAGGATTCACAAGCGCATCGTGAATAAGAGACAAGATTTTCTGCACAAACTGACCACTTCCCTTGTGCGCGAGTATGAAATAATCGGGACTGAACACCTGAAGCCCGACAATATGCGCAAGAATAGAAGGCTGGCGCTCAGCATCTCAGATGCTGGATGGGGTGAATTCATAAGGCAGTTGGAGTACAAAGCTGCTTGGTATGGGAGACTCGTAAGTAAAGTTTCCCCATATTTTCCTTCTAGTCAACTCTGTCACGACTGCGGATTCAAAAACCCCGAAGTGAAAAATCTCGCGGTAAGGACTTGGACGTGCCCAAACTGTGGCGAGACACACGACAGAGATGAAAATGCGGCGCTGAATATCAGGAGAGAGGCGTTGGTGGCTGCGGGAATCTCAGACACCTTAAACGCTCATGGAGGCTATGTCAGACCTGCTTCGGCGGGCAATGGTCTGCGAAGTGAGAATCACGCGACTTTAGTCGTGTGAGGTTCAAGAAGAGATCTCTGAACACCCggccggcatggtcccagcctcctcgctggcgccggctgggcaacatgcttcggcatggcgaatgggacccgtagctcagttggttagagcgttggtcttatgagccgaaggtcgcgggttcgagccccgccggaagca

**Supplementary sequence 6**

**TRV-DTU.1 sequence for Nicotiana benthamiana genome editing**

NLS-DTU.1 TnpB-HH-reRNA (116 bp)-Spacer)-HDV-tRNA^ile^

ATGGCCCCAAAGAAGAAGCGCAAGGTCATACGCtatAAGGCATTTGTGGTTCGGCTCTACCCGAATGCGGCACAAACTGAGCTGATAAACCGAACACTGGGGTCTGCACGATTTGTATACAATCATTTCCTTGCTCGAAGAATCGCTGCGTACAAGGAAAGCGGTAAGGGTCTTACGTACGGGCAAACTTCTAGTGAGCTTACTCTGCTGAAGCAAGCAGAGGAAACCTCCTGGCTGTCAGAAGTGGATAAATTCGCGTTGCAGAATAGCTTGAAAAACCTTGAAACGGCCTATAAAAACTTCTTTAGAACAGTTAAGCAGTCAGGAAAAAAGGTAGGTTTCCCTCGGTTCaaaAAGAAGCGCACGGGGGAGTCATACCGCACGCAGTTTACGAACAACAACATTCAGATAGGAGAGGGGAGGCTTAAACTGCCGAAACTGGGGTGGGTCAAAACTAAGGGTCAGCAGGATATCCAAGGCAAAATATTGAATGTTACTGTACGCCGCATCCATGAAGGCCATTATGAAGCTTCAGTGCTGTGCGAGGTCGAAATCCCTTACCTCCCAGCTGCTCCGAAATTCGCGGCAGGAGTAGACCTGGGGATTAAAGACTTTGCGATCGTCACCGATGGTGTGCGATTTAAGCATGAGCAAAATCCTAAATACTATCGGAGTACGCTTAAGCGAATTCGGAAGGCCCAACAGACTCTGTCCAGGAGAAAGAAAGGAAGCGCTAGATATGGCAAGGCTAAGACTAAGCTGGCAAGGATTCACAAGCGCATCGTGAATAAGAGACAAGATTTTCTGCACAAACTGACCACTTCCCTTGTGCGCGAGTATGAAATAATCGGGACTGAACACCTGAAGCCCGACAATATGCGCAAGAATAGAAGGCTGGCGCTCAGCATCTCAGATGCTGGATGGGGTGAATTCATAAGGCAGTTGGAGTACAAAGCTGCTTGGTATGGGAGACTCGTAAGTAAAGTTTCCCCATATTTTCCTTCTAGTCAACTCTGTCACGACTGCGGATTCAAAAACCCCGAAGTGAAAAATCTCGCGGTAAGGACTTGGACGTGCCCAAACTGTGGCGAGACACACGACAGAGATGAAAATGCGGCGCTGAATATCAGGAGAGAGGCGTTGGTGGCTGCGGGGATTTCAGATACCCTTAACGCGCACGGCGGTTACGTGCGACCCGCTTCCGCCGGTAACGGCCTTAGGAGTGAAAACCACGCTACTCTCGTAGTGTAGggtaccttcttgaatcctgatgagtccgtgaggacgaaacgagctagctcgtcGGTGGCTGCGGGAATCTCAGACACCTTAAACGCTCATGGAGGCTATGTCAGACCTGCTTCGGCGGGCAATGGTCTGCGAAGTGAGAATCACGCGACTTTAGTCGTGTGAGGTTCAAGCTACAATGAAGGAACTAGCggccggcatggtcccagcctcctcgctggcgccggctgggcaacatgcttcggcatggcgaatgggacGAGCTCCCGTAGCTCAGTTGGTTAGAGCGTTGGTCTTATGAGCCGAAGGTCGCGGGTTCGAGCCCCGCCGGAAGCA

**Supplementary sequence 7**

**VS-STU.2 sequence for Nicotiana benthamiana genome editing**

NLS-STU.2 TnpB (reRNA (224 bp unoptimized)-Stop Codon (TGA)- GGTTCAA-Spacer)-HDV-NbPDS-tRNA^ile^

ATGGCCCCAAAGAAGAAGCGCAAGGTCATACGCtatAAGGCATTTGTGGTTCGGCTCTACCCGAATGCGGCACAAACTGAGCTGATAAACCGAACACTGGGGTCTGCACGATTTGTATACAATCATTTCCTTGCTCGAAGAATCGCTGCGTACAAGGAAAGCGGTAAGGGTCTTACGTACGGGCAAACTTCTAGTGAGCTTACTCTGCTGAAGCAAGCAGAGGAAACCTCCTGGCTGTCAGAAGTGGATAAATTCGCGTTGCAGAATAGCTTGAAAAACCTTGAAACGGCCTATAAAAACTTCTTTAGAACAGTTAAGCAGTCAGGAAAAAAGGTAGGTTTCCCTCGGTTCaaaAAGAAGCGCACGGGGGAGTCATACCGCACGCAGTTTACGAACAACAACATTCAGATAGGAGAGGGGAGGCTTAAACTGCCGAAACTGGGGTGGGTCAAAACTAAGGGTCAGCAGGATATCCAAGGCAAAATATTGAATGTTACTGTACGCCGCATCCATGAAGGCCATTATGAAGCTTCAGTGCTGTGCGAGGTCGAAATCCCTTACCTCCCAGCTGCTCCGAAATTCGCGGCAGGAGTAGACCTGGGGATTAAAGACTTTGCGATCGTCACCGATGGTGTGCGATTTAAGCATGAGCAAAATCCTAAATACTATCGGAGTACGCTTAAGCGAATTCGGAAGGCCCAACAGACTCTGTCCAGGAGAAAGAAAGGAAGCGCTAGATATGGCAAGGCTAAGACTAAGCTGGCAAGGATTCACAAGCGCATCGTGAATAAGAGACAAGATTTTCTGCACAAACTGACCACTTCCCTTGTGCGCGAGTATGAAATAATCGGGACTGAACACCTGAAGCCCGACAATATGCGCAAGAATAGAAGGCTGGCGCTCAGCATCTCAGATGCTGGATGGGGTGAATTCATAAGGCAGTTGGAGTACAAAGCTGCTTGGTATGGGAGACTCGTAAGTAAAGTTTCCCCATATTTTCCTTCTAGTCAACTCTGTCACGACTGCGGATTCAAGAATCCCGAAGTGAAGAATCTTGCCGTCCGTACATGGACTTGCCCGAACTGTGGGGAAACCCATGACCGAGACGAGAACGCTGCGCTGAACATTCGGCGTGAAGCGTTGGTGGCTGCGGGAATCTCAGACACCTTAAACGCTCATGGAGGCTATGTCAGACCTGCTTCGGCGGGCAATGGTCTGCGAAGTGAGAATCACGCGACTTTAGTCGTGTGAGGTTCAAGAAGAGATCTCTGAACACCCggccggcatggtcccagcctcctcgctggcgccggctgggcaacatgcttcggcatggcgaatgggacCACAGGAACTCCCACTAGCTTCTCCAACTTTTGGAAATATGGGATCTCTTTCCAGTCTTCAGGCAAAAGAAGCTTCAAGATATCCACTAATTGGATAATCAAAAGTCAGCAAAAAACTATAAATTCATATTAAAATGTGAAAACTGGAAGCGTCATACTAGTGTAATGGATATTATACCTGGAGTGGCAAACACAAAAGCATCTCCTTTAATTGTACTGCCATTATTCAGTATAAAACATTTGACACTTCCATCCTCATTCAGCTCGATCTTTTTTATTCGTGAGTTTAGTCTGACTTGGccgtagctcagttggttagagcgttggtcttatgagccgaaggtcgcgggttcgagccccgccggaagca

**Supplementary sequence 7**

**VS-STU.3 sequence for Nicotiana benthamiana genome editing**

NLS-STU.3 TnpB (reRNA (224 bp unoptimized)-Stop Codon (TGA)- GGTTCAA-Spacer)-HDV-NbPDS-tRNA^ile^

ATGGCCCCAAAGAAGAAGCGCAAGGTCATACGCtatAAGGCATTTGTGGTTCGGCTCTACCCGAATGCGGCACAAACTGAGCTGATAAACCGAACACTGGGGTCTGCACGATTTGTATACAATCATTTCCTTGCTCGAAGAATCGCTGCGTACAAGGAAAGCGGTAAGGGTCTTACGTACGGGCAAACTTCTAGTGAGCTTACTCTGCTGAAGCAAGCAGAGGAAACCTCCTGGCTGTCAGAAGTGGATAAATTCGCGTTGCAGAATAGCTTGAAAAACCTTGAAACGGCCTATAAAAACTTCTTTAGAACAGTTAAGCAGTCAGGAAAAAAGGTAGGTTTCCCTCGGTTCaaaAAGAAGCGCACGGGGGAGTCATACCGCACGCAGTTTACGAACAACAACATTCAGATAGGAGAGGGGAGGCTTAAACTGCCGAAACTGGGGTGGGTCAAAACTAAGGGTCAGCAGGATATCCAAGGCAAAATATTGAATGTTACTGTACGCCGCATCCATGAAGGCCATTATGAAGCTTCAGTGCTGTGCGAGGTCGAAATCCCTTACCTCCCAGCTGCTCCGAAATTCGCGGCAGGAGTAGACCTGGGGATTAAAGACTTTGCGATCGTCACCGATGGTGTGCGATTTAAGCATGAGCAAAATCCTAAATACTATCGGAGTACGCTTAAGCGAATTCGGAAGGCCCAACAGACTCTGTCCAGGAGAAAGAAAGGAAGCGCTAGATATGGCAAGGCTAAGACTAAGCTGGCAAGGATTCACAAGCGCATCGTGAATAAGAGACAAGATTTTCTGCACAAACTGACCACTTCCCTTGTGCGCGAGTATGAAATAATCGGGACTGAACACCTGAAGCCCGACAATATGCGCAAGAATAGAAGGCTGGCGCTCAGCATCTCAGATGCTGGATGGGGTGAATTCATAAGGCAGTTGGAGTACAAAGCTGCTTGGTATGGGAGACTCGTAAGTAAAGTTTCCCCATATTTTCCTTCTAGTCAACTCTGTCACGACTGCGGATTCAAAAACCCCGAAGTGAAAAATCTCGCGGTAAGGACTTGGACGTGCCCAAACTGTGGCGAGACACACGACAGAGATGAAAATGCGGCGCTGAATATCAGGAGAGAGGCGTTGGTGGCTGCGGGAATCTCAGACACCTTAAACGCTCATGGAGGCTATGTCAGACCTGCTTCGGCGGGCAATGGTCTGCGAAGTGAGAATCACGCGACTTTAGTCGTGTGAGGTTCAAGAAGAGATCTCTGAACACCCggccggcatggtcccagcctcctcgctggcgccggctgggcaacatgcttcggcatggcgaatgggacCACAGGAACTCCCACTAGCTTCTCCAACTTTTGGAAATATGGGATCTCTTTCCAGTCTTCAGGCAAAAGAAGCTTCAAGATATCCACTAATTGGATAATCAAAAGTCAGCAAAAAACTATAAATTCATATTAAAATGTGAAAACTGGAAGCGTCATACTAGTGTAATGGATATTATACCTGGAGTGGCAAACACAAAAGCATCTCCTTTAATTGTACTGCCATTATTCAGTATAAAACATTTGACACTTCCATCCTCATTCAGCTCGATCTTTTTTATTCGTGAGTTTAGTCTGACTTGGccgtagctcagttggttagagcgttggtcttatgagccgaaggtcgcgggttcgagccccgccggaagca

**Supplementary sequence 8**

**VS-DTU.1 sequence for Nicotiana benthamiana genome editing**

NLS-DTU.1 TnpB-HH-reRNA (116 bp)-Spacer)-HDV-tRNA^ile^

ATGGCCCCAAAGAAGAAGCGCAAGGTCATACGCtatAAGGCATTTGTGGTTCGGCTCTACCCGAATGCGGCACAAACTGAGCTGATAAACCGAACACTGGGGTCTGCACGATTTGTATACAATCATTTCCTTGCTCGAAGAATCGCTGCGTACAAGGAAAGCGGTAAGGGTCTTACGTACGGGCAAACTTCTAGTGAGCTTACTCTGCTGAAGCAAGCAGAGGAAACCTCCTGGCTGTCAGAAGTGGATAAATTCGCGTTGCAGAATAGCTTGAAAAACCTTGAAACGGCCTATAAAAACTTCTTTAGAACAGTTAAGCAGTCAGGAAAAAAGGTAGGTTTCCCTCGGTTCaaaAAGAAGCGCACGGGGGAGTCATACCGCACGCAGTTTACGAACAACAACATTCAGATAGGAGAGGGGAGGCTTAAACTGCCGAAACTGGGGTGGGTCAAAACTAAGGGTCAGCAGGATATCCAAGGCAAAATATTGAATGTTACTGTACGCCGCATCCATGAAGGCCATTATGAAGCTTCAGTGCTGTGCGAGGTCGAAATCCCTTACCTCCCAGCTGCTCCGAAATTCGCGGCAGGAGTAGACCTGGGGATTAAAGACTTTGCGATCGTCACCGATGGTGTGCGATTTAAGCATGAGCAAAATCCTAAATACTATCGGAGTACGCTTAAGCGAATTCGGAAGGCCCAACAGACTCTGTCCAGGAGAAAGAAAGGAAGCGCTAGATATGGCAAGGCTAAGACTAAGCTGGCAAGGATTCACAAGCGCATCGTGAATAAGAGACAAGATTTTCTGCACAAACTGACCACTTCCCTTGTGCGCGAGTATGAAATAATCGGGACTGAACACCTGAAGCCCGACAATATGCGCAAGAATAGAAGGCTGGCGCTCAGCATCTCAGATGCTGGATGGGGTGAATTCATAAGGCAGTTGGAGTACAAAGCTGCTTGGTATGGGAGACTCGTAAGTAAAGTTTCCCCATATTTTCCTTCTAGTCAACTCTGTCACGACTGCGGATTCAAAAACCCCGAAGTGAAAAATCTCGCGGTAAGGACTTGGACGTGCCCAAACTGTGGCGAGACACACGACAGAGATGAAAATGCGGCGCTGAATATCAGGAGAGAGGCGTTGGTGGCTGCGGGGATTTCAGATACCCTTAACGCGCACGGCGGTTACGTGCGACCCGCTTCCGCCGGTAACGGCCTTAGGAGTGAAAACCACGCTACTCTCGTAGTGTAGggtaccttcttgaatcctgatgagtccgtgaggacgaaacgagctagctcgtcGGTGGCTGCGGGAATCTCAGACACCTTAAACGCTCATGGAGGCTATGTCAGACCTGCTTCGGCGGGCAATGGTCTGCGAAGTGAGAATCACGCGACTTTAGTCGTGTGAGGTTCAAGCTACAATGAAGGAACTAGCggccggcatggtcccagcctcctcgctggcgccggctgggcaacatgcttcggcatggcgaatgggacCACAGGAACTCCCACTAGCTTCTCCAACTTTTGGAAATATGGGATCTCTTTCCAGTCTTCAGGCAAAAGAAGCTTCAAGATATCCACTAATTGGATAATCAAAAGTCAGCAAAAAACTATAAATTCATATTAAAATGTGAAAACTGGAAGCGTCATACTAGTGTAATGGATATTATACCTGGAGTGGCAAACACAAAAGCATCTCCTTTAATTGTACTGCCATTATTCAGTATAAAACATTTGACACTTCCATCCTCATTCAGCTCGATCTTTTTTATTCGTGAGTTTAGTCTGACTTGGCCGTAGCTCAGTTGGTTAGAGCGTTGGTCTTATGAGCCGAAGGTCGCGGGTTCGAGCCCCGCCGGAAGCA
